# Single Cell Enhancer Activity Maps Neuronal Lineages in Embryonic Mouse Basal Ganglia

**DOI:** 10.1101/2021.01.11.426285

**Authors:** Linda Su-Feher, Anna N. Rubin, Shanni N. Silberberg, Rinaldo Catta-Preta, Kenneth J. Lim, Iva Zdilar, Christopher S. McGinnis, Gabriel L. McKinsey, Thomas E. Rubino, Michael Hawrylycz, Carol Thompson, Zev J. Gartner, Luis Puelles, Hongkui Zeng, John L. R. Rubenstein, Alex S. Nord

**Affiliations:** Department of Psychiatry and Behavioral Sciences, University of California, Davis, Davis, California, USA; Department of Neurobiology, Physiology and Behavior, University of California, Davis, Davis, California, USA; Nina Ireland Laboratory of Developmental Neurobiology, Department of Psychiatry, University of California, San Francisco Medical School, San Francisco, California, USA; Department of Pharmaceutical Chemistry, University of California, San Francisco, San Francisco, CA, USA; Department of Pediatrics, University of California, San Francisco, San Francisco, USA; Allen Institute for Brain Science, Seattle, WA, USA; Helen Diller Family Comprehensive Cancer Center, San Francisco, CA, USA; Chan Zuckerberg BioHub, University of California San Francisco, San Francisco, CA, USA; Center for Cellular Construction, University of California San Francisco, San Francisco, CA, USA.; Department of Human Anatomy and Psychobiology and IMIB-Arrixaca Institute, University of Murcia, Spain; Department of Genetics, Blavatnik Institute, Harvard Medical School, Boston, Massachusetts, USA

**Author notes:** These authors contributed equally (Linda Su-Feher, Anna N. Rubin). Correspondence to Alex S. Nord and John L. R. Rubenstein.

## Abstract

Enhancers integrate transcription factor signaling pathways that drive cell fate specification in the developing brain. We used single cell RNA-sequencing (scRNA-seq) to capture enhancer activity at single cell resolution and delineate specification of cells labeled by enhancers in mouse medial, lateral, and caudal ganglionic eminences (MGE, LGE, and CGE) at embryonic day (E)11.5. We combine enhancer-based reporter labeling with single-cell transcriptional readout to characterize enhancer activity and define cell populations in vivo. Seven enhancers had diverse activities in specific progenitor and neuronal populations within the GEs. We then applied enhancer-based labeling, scRNA-seq, and analysis of in situ hybridization (ISH) data to distinguish subtypes of MGE-derived GABAergic and cholinergic projection neurons and interneurons. This work demonstrates how the power of scRNA-seq can be extended by enhancer-based labelling and leveraging ISH data and reveals novel lineage specification paths underlying patterning of developing mouse brain.

## Introduction

During brain development, transcriptional programs governed by the genomic interplay of transcription factors and cis-regulatory enhancer and promoter sequences drive the proliferation and specification of neuronal and glial lineages (Beccari et al., 2013; Nord, 2015). An understanding of this regulatory symphony in the telencephalon has been derived via decades of genetic dissection of transcription factor signaling (Kessaris et al., 2014; Lim et al., 2018; Long et al., 2009), more recently extended via genomic approaches (Lindtner et al., 2019; Sandberg et al., 2016), and is now undergoing a revolution via application of single cell RNA-sequencing (scRNA-seq). scRNA-seq has produced fine-scale elucidation of cell types in the mammalian brain (Zeisel et al., 2018); however, major challenges remain towards understanding the dynamics of cell state and identity that occur in the context of neurodevelopment.

The embryonic basal ganglia (BG) include spatially distinct proliferative zones of the ganglionic eminences (GEs), which include the medial, lateral, and caudal ganglionic eminences (MGE, LGE, and CGE) (J.L.R. and Campbell, 2020). Progenitor cells in the ventricular (VZ) and subventricular (SVZ) domains in the GEs give rise to many neuronal classes. Neuron types that originate in embryonic BG include GABAergic projection neurons and cholinergic neurons (Fragkouli et al., 2009) that form the ventral pallidum, globus pallidus (Flandin et al., 2010; Nóbrega-Pereira et al., 2010), and striatal structures (J.L.R. and Campbell, 2020) that make up the mature BG. In addition, the GEs generate interneurons that populate the striatum, cortex, olfactory bulb, and other brain regions (Anderson et al., 1997; Batista-Brito et al., 2020; Lim et al., 2018; Marín et al., 2000). Building on bulk transcriptomics and in situ hybridization studies (ISH), scRNA-seq has been applied to embryonic mouse BG, revealing generalized progenitor populations and early born GABAergic lineages, with a focus on cortical interneuron (CIN) specification (Mayer et al., 2018; Mi et al., 2018). While CINs are one major output of embryonic BG, single cell characterization of GABAergic and cholinergic as well as early born CIN lineages and that arise in the BG remains largely unexplored. Resolving the early stages of BG neurogenesis via scRNA-seq and ISH has been limited by major barriers: paucity of region- and lineage-specific single gene markers, similarity of early transcriptional programs, spatial mixing of progenitors within germinal zones and immature cell types in the MZ, and regional organization of BG neurogenesis that has been poorly captured by unguided scRNA-seq analysis.

Fate mapping via reporter labeling has provided critical insights into the origins of neuronal cell populations (Batista-Brito et al., 2020). Notably, enhancers drive highly specific transcription pattens, including in the developing telencephalon (Visel et al., 2013), thus offering exciting possibilities for cell-type specific labeling and genetic manipulation. We previously demonstrated the utility of enhancer-driven transgenic reporter mouse lines for fate mapping and genetic manipulation of neuronal populations originating in embryonic BG and cortex (Pattabiraman et al., 2014; Silberberg et al., 2016). We generated transgenic mice harboring evolutionarily conserved enhancer sequences that drive expression of CreER^T2^ and GFP. These developmental enhancers exhibited spatiotemporal activity across expression domains within the embryonic BG and mark early cell populations prior to terminal cell fate commitment. Enhancers differentially labeled cell populations that spatially intermingle and alternatively marked regionally distinct mitotic and postmitotic populations during development. Fate mapping with these enhancer-driven CreER^T2^-GFP mice demonstrated that developmental lineages marked by transient enhancer activity produce varied mature neuron populations within and across these enhancers. Importantly, beyond their use in understanding neuronal lineages, these enhancer-driven reporter lines offer the opportunity for function-based analysis of dynamic in vivo enhancer activity, a missing feature from studies modeling enhancer activity via epigenomic approaches. More broadly, enhancer-based cell labeling is emerging as a powerful tool for cell-type identification, enrichment, and modulation in neuroscience and other areas, yet little remains known about sensitivity and specificity of enhancer-driven reporter expression at single cell resolution.

In this study, we apply the novel strategy of pairing enhancer-based transgenic reporter mouse lines with scRNA-seq to define specific enhancer-labeled lineages at single cell resolution in early embryonic BG. These experiments reveal functionally defined distinct enhancer activities across scRNA-seq-defined cell states and lineages. Next, we focused on the MGE and integrated ISH mapping of transcript expression from the Allen Developing Mouse Brain Atlas (ABA) (Lein et al., 2007) to provide a higher resolution anatomical definition of lineages identified by enhancer-labeling and scRNA-seq. Our study identified proliferative and postmitotic cells that are distinctly labeled by enhancers active in MGE, LGE, and CGE and revealed novel specification paths for enhancer-labeled and spatially defined populations of early BG-derived neuronal lineages.

## Results

### Comparative activity of seven enhancers in E11.5 BG via scRNA-seq

We profiled enhancer-labeled cell populations from day (E)11.5 MGE, LGE, or CGE across seven subpallial enhancer transgenic mouse lines (Silberberg et al., 2016) (Figure 1A). The selected transgenic lines express GFP and CreER^T2^ with enhancer-driven divergent patterns in the ventricular (VZ), subventricular (SVZ), and mantle (MZ) zones of the GEs. These enhancers are putatively associated with developmentally expressed genes and have restricted regional activity within the GEs at E11.5, summarized in Figure 1B. The objective of these experiments was threefold. First, to establish the utility and sensitivity to detect enhancer-driven reporter expression via scRNA-seq. Second, to define and compare representative enhancer activities and enhancer-labeled progenitors and early neuronal populations across MGE (enhancers *hs1538*, *hs1056*, *hs799*, and *hs192*), LGEs (*hs841* and *hs599*), and CGE (*hs841* and *hs953*) at E11.5. Third, to resolve fine-scale differences among cells labeled by regionally distinct MGE progenitor-associated enhancers (*hs1538* and *hs1056*) and early neuronal enhancers that differentially label emerging lineages (*hs799* and *hs192*).

**Figure 1:**
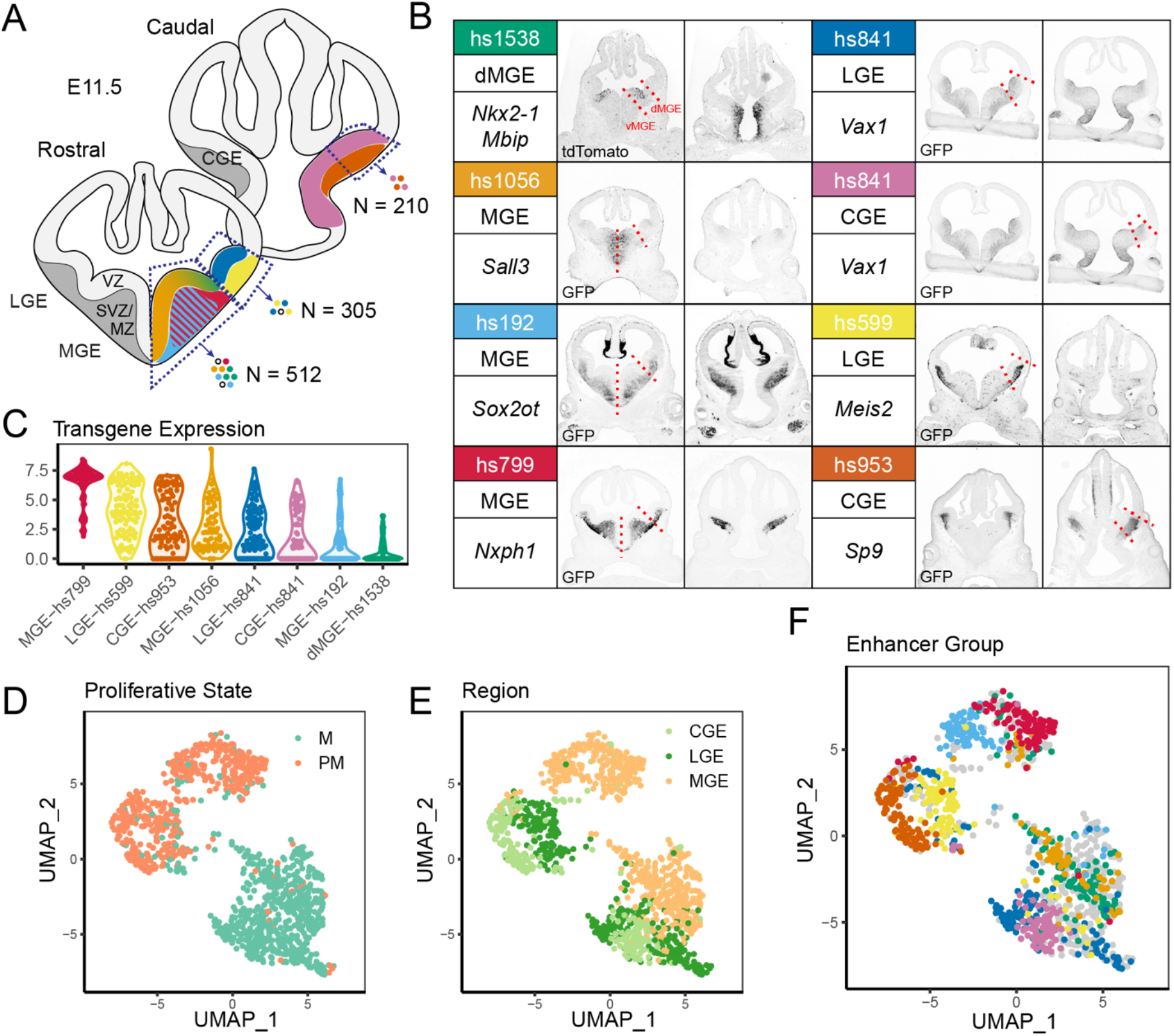
Profiling enhancer-labeled single cells in E11.5 basal ganglia. (**A**) Schematic of dissection, with colors representing the activity of seven transgenic enhancer reporters characterized using C1 scRNA-seq. CGE: caudal ganglionic eminence; LGE: lateral ganglionic eminence; MGE: medial ganglionic eminence; VZ: ventricular zone; SVZ: subventricular zone; MZ: mantle zone. (**B**) Summary of the seven enhancers profiled, including dissection region, putative gene regulatory target, and representative GFP immunohistochemistry (IHC) imaging of enhancer transgenic reporters at E11.5, depicting activity within the ganglionic eminences. Red lines indicate microdissection boundaries. GFP IHC images are adapted from (Silberberg et al., 2016). (**C**) Violin plot of normalized transgene expression by enhancer group. (**D**) Visualization of single cells by UMAP, colored by mitotic state (green: M, mitotic; orange: PM, postmitotic). (**E**) Visualization of single cells by UMAP, colored by region of dissection (light orange: MGE; light green: CGE; dark green: LGE). (**F**) Visualization of single cells by UMAP, colored by transgenic enhancer grouping. Colors correspond to header colors in (B). Enhancer-negative cells are depicted in grey.

Using the seven transgenic lines, we performed targeted BG microdissection and preparation of reporter-positive and ungated single cells. Single cell suspensions were either first segregated for transgene expression through fluorescence activated cell sorting (FACS, Figure S1) or passed directly to the Fluidigm C1 system for capture and amplification of the transcriptomes of individual cells. The regional dissections included: for *hs599* the LGE; for *hs953* the CGE; for *hs1538* the dorsal (d)MGE; and for *hs1056, hs192*, and *hs799*, the MGE. For one enhancer, *hs841*, we independently dissected the LGE and CGE. For *hs1538*, we used CreER^T2^-driven tdTomato signal via cross to Ai14 reporter mice (Madisen et al., 2010) for gating due to low GFP signal. For details regarding sample preparation, see Table S1. After sequencing and quality control (Figure S2A-L), 1027 cells were included for analysis, with ∼594,000 reads and ∼5,140 genes per cell on average.

Our first objective was to demonstrate the feasibility of single cell enhancer activity mapping by establishing whether enhancer-driven transgene expression could be mapped to single cells via scRNA-seq. We used a combination of FACS^+^ gating and transgene (CreER^T2^-IRES-GFP or tdTomato) RNA expression to assign cells as enhancer-positive or negative (“None”). 315 cells were unsorted or FACS^-^ and transgene negative; 712 cells were FACS^+^ and/or expressed non-zero transgene. Based on tissue dissection and enhancer line, enhancer-positive cells were assigned to one of eight categories: MGE-*hs1056*, MGE-*hs1538*, MGE-*hs192*, MGE-*hs799*, LGE-*hs599*, CGE-*hs953*, LGE-*hs841*, and CGE-*hs841*. Enhancer-labeled cells generally exhibited expression of presumed target genes with some exceptions. Enhancer-driven reporter transcripts were detectable at single cell resolution across all enhancers, with variation in presence and transcript level captured via scRNA-seq (Figure 1C). Five of the seven enhancers exhibited strong concordance between reporter protein GFP^+^ gating and transgene transcript detected via scRNA-seq (Figure S2M). The other two lines, dMGE-*hs1538* and MGE-*hs192*, had weaker sensitivity, with 30-40% of FACS reporter-positive cells having detectable transgene transcript. Nonetheless, even for enhancers with weaker transcriptional activity, reporter transgene was reliably detected via scRNA-seq in a substantial fraction of FACS-determined reporter-positive cells, demonstrating the overall utility of this approach for function-based scRNA-seq enhancer activity profiling.

### TF expression organizes scRNA-seq data by proliferative state and BG region

Using highly variable genes in scRNA-seq analysis is a common approach for feature selection (Butler et al., 2018); however, this method did not adequately separate regional and cell state identity in our data (Figure S3). As an alternative, we used a transcription factor (TF)-curated approach, with the rationale that TFs drive lineage specification and cell identity. We rooted this analysis using 689 TFs profiled for RNA ISH patterns at E11.5 and E13.5 in the Allen Developing Mouse Brain Atlas (ABA) (Lein et al., 2007) (Table S2, Figure S4A-B). 455 of these TFs were expressed in our E11.5 scRNA-seq data, of which 292 (64.2%) had detectable ISH expression in the BG (Figure S4C-E). We used these 455 TFs to define scRNA-seq cell identity and for visual representation via UMAP plots (Figure 1D-F). Using this TF-curated approach, proliferative state and regional origin were the primary aspects of scRNA-seq variation (Figures 1D-E). We compared TF-curated analysis to results using highly variable genes (Figures S3A-C), before and after performing regression analysis to reduce the influence of cell cycle phase (Figures S3D-F). Excluding non-TFs reduced the contribution of cell cycle phase and confounding sources of variation (e.g. sequencing batch) to cell clustering and improved separation by GE origin (Figure S3G-I).

### Enhancers label cells with specific regional identities and developmental trajectories

Having shown that scRNA-seq can reliably identify enhancer-positive cells and that our TF-curated approach enables separation across regional origin and proliferative states, we next modeled transcriptional differences across enhancer-positive cells. First, we modeled transitional cell states via diffusion mapping. Second, we examined differences across cluster-based transcriptional identities. We identified two major diffusion components (DC) corresponding to proliferative state (DC1) and MGE from LGE/CGE origin (DC2) (Figure 2A-F). DC1 captured the stem cell, proneural, and neurogenic transition, with genes such as *Hes1*, *Ccnd2*, *Gadd45g*, and *Slc34a2* marking cells at various stages of this transition (Figure 2G). Lower values of DC2 were associated with MGE identity, marked by expression such MGE-specific genes such as *Nkx2-1*, *Lhx6*, and *Lhx8* (Flames et al., 2007) (Figure 2G). Higher values of DC2 were associated with LGE or CGE identity (Figure 2D,F). DC2 diversity was driven by expression of region-defining TFs such as *Nkx2-1* in the MGE and *Pax6* in the CGE and LGE (Figure 2G,I).

**Figure 2:**
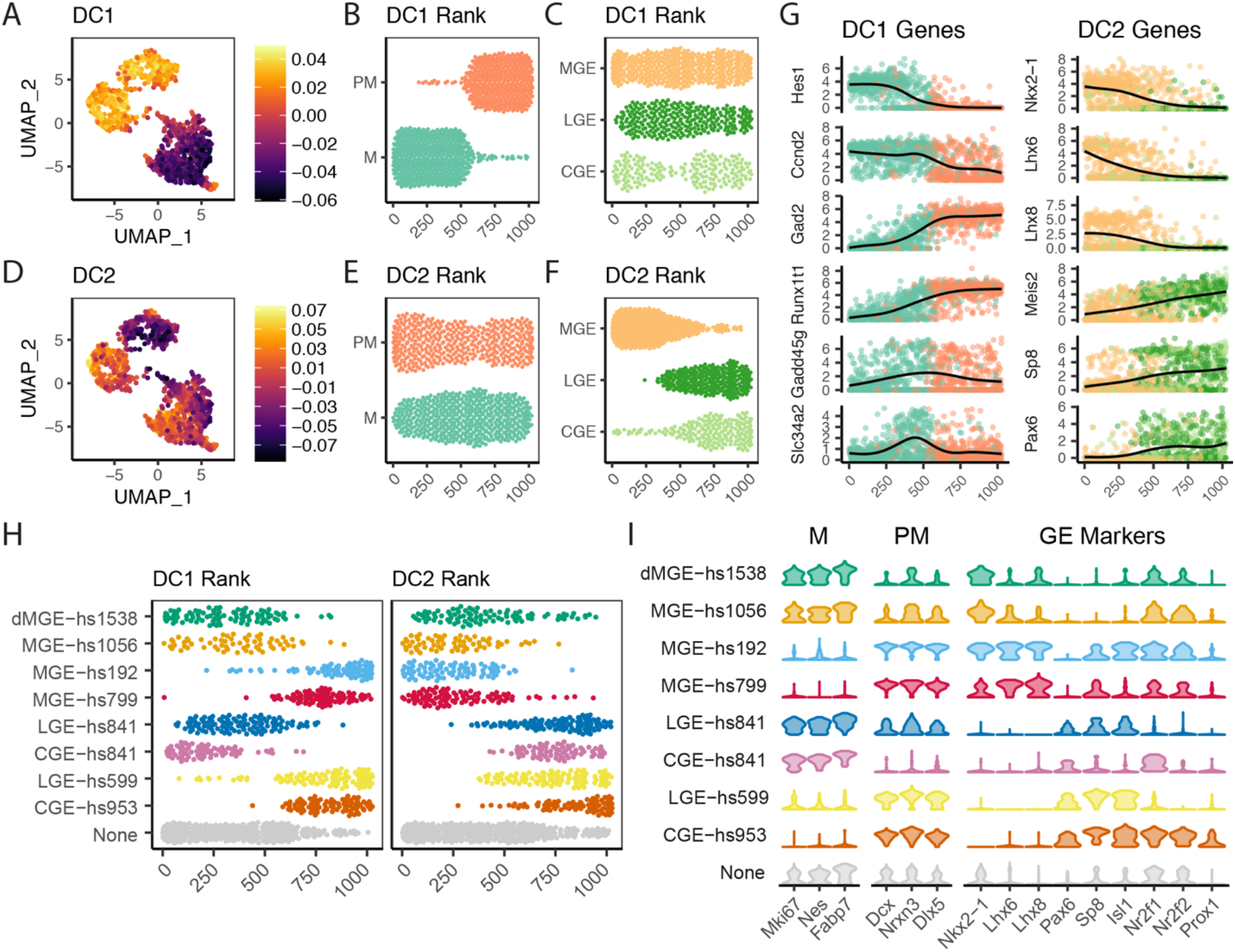
Diffusion mapping reveals progenitor state and regional identity gradients. (**A,D**) UMAP colored by diffusion component value (DC)1 (A) or DC2 (D). (**B,E**) Cells ranked by DC1 (B) or DC2 (E) value, separated and colored by mitotic state. (**C,F**) Cells ranked by DC1 (B) or DC2 (E) value, separated and colored by ganglionic eminence. (**G**) Relative expression of differentially expressed genes across DC1 (*left*) or DC2 (*right*). Cells on x-axis are ordered by DC1 or DC2 rank. Line represents generalized additive model (gam) line. (**H**) Cells ranked by DC1 (*left*) or DC2 (*right*) rank, separated and colored by enhancer group. (**I**) Violin plots of relative expression by enhancer group for marker genes associated with mitotic identity (M), postmitotic identity (PM), and various markers with ganglionic eminence-associated expression.

The strongest separation of LGE and CGE identity are the caudal-biased TFs *Nr2f1* and *Nr2f2* (Hu et al., 2017) (Figure 2I). DC1 and DC2 values distinguished cells labeled by different enhancers and indicate that these developmental enhancers are active across maturation states within the GEs (Figure 2H). VZ-associated enhancers dMGE-*hs1538*, MGE-*hs1056*, and CGE- and LGE-*hs841* labeled cells across the proliferative zone of DC1, indicating enhancer activity across multiple maturation states (Figure 2H). In contrast, SVZ/MZ-associated enhancers MGE-*hs192*, MGE-*hs799*, LGE-*hs599*, and CGE-*hs953* labeled cells across the postmitotic zone of DC1, indicating these enhancers are active across neuronal maturation (Figure 2H).

We next performed clustering using TF-curated scRNA-seq expression, identifying 12 cell clusters that separated by proliferative state and regional or cell-type identity (Figure 3A, Table S3). We further used random forest classification to define informative transcripts that discriminate cells labelled by specific enhancers (Table S4). Cells labeled by enhancers dMGE-*hs1538*, MGE-*hs1056*, CGE-*hs841* and LGE-*hs841* primarily grouped into mitotic clusters (cl)-1, cl-2, cl-3, cl-8, and cl-9, further separated by regional identity (MGE versus non-MGE). Within these regional boundaries, VZ/SVZ-associated enhancers split across multiple clusters (Figure 3B), paralleling diffusion mapping results suggesting mitotic enhancers label multiple proliferative states. Compared to enhancers with progenitor activity, enhancers active in postmitotic cells (MGE-*hs192*, MGE-*hs799*, LGE-*hs599*, and CGE-*hs953*) were biased toward specific cell type clusters within broader regional identities (Figure 3B). To characterize cell types that were differentially labeled by these enhancers, we performed differential gene expression analysis using the full transcriptome of 17,015 expressed genes to identify differentially expressed (DE) genes for each TF-defined cluster (Figure 3C-E).

**Figure 3:**
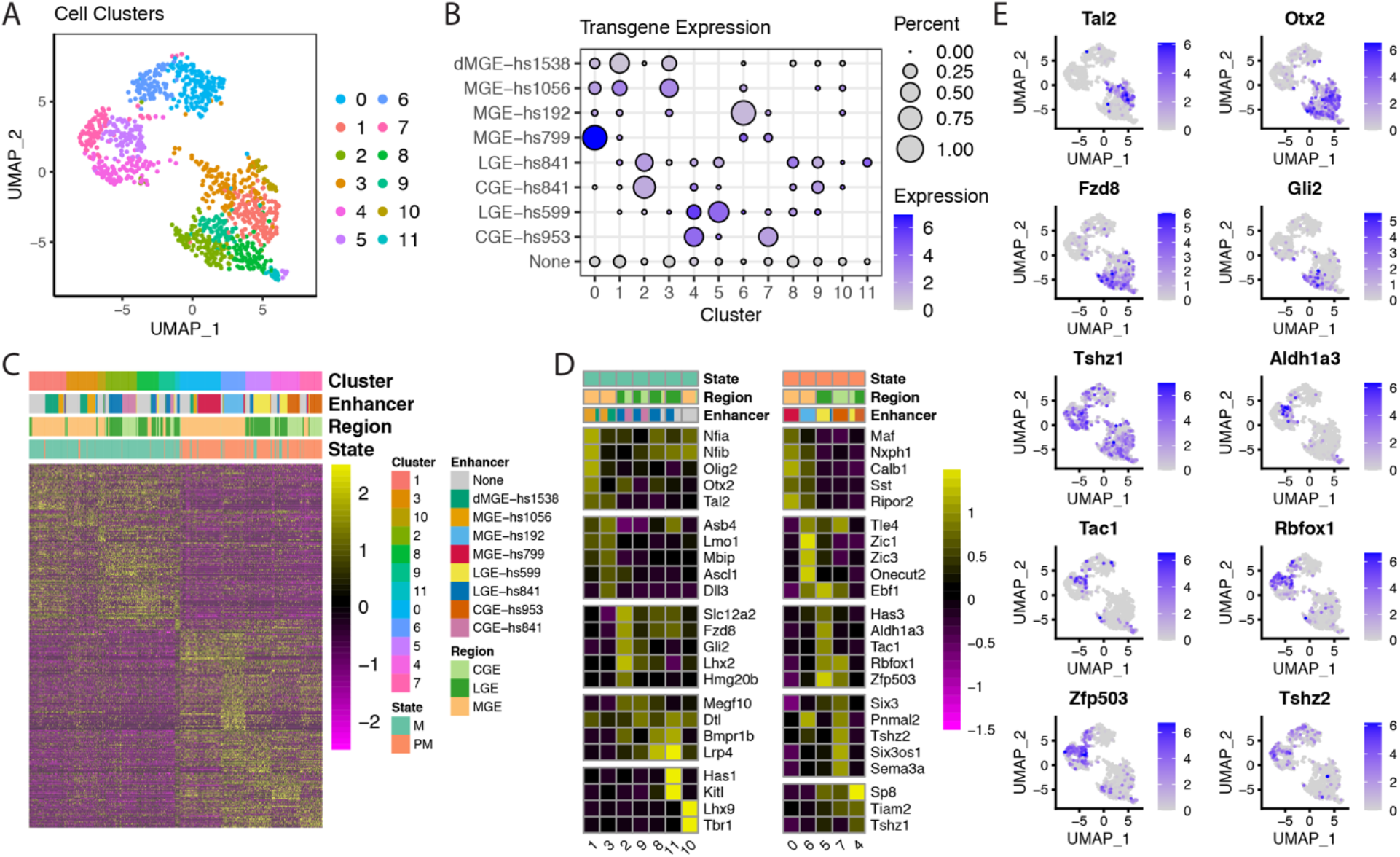
Enhancers label distinct transcriptional signatures. (**A**) UMAP colored by TF-defined cell cluster. (**B**) Dot plot of relative transgene expression and representation of each enhancer across clusters. Size of dot represents percent of enhancer corresponding to a specific cluster. Color gradient represents mean normalized transgene expression for each enhancer by cluster. (**C**) Ordered heatmap of differentially expressed genes, in order of cluster, enhancer, regional origin, and proliferative state. Color gradient represents Z-score of normalized expression. (**D**) Representative differentially expressed genes by cluster. Color represents the mean Z-score of normalized expression across clusters. Each column color bar represents the proportion representation of proliferative state, region, or enhancer group. (**E**) UMAP plots of representative genes from (D), colored by relative expression.

Proliferative clusters cl-1, 3, 2, and 9 encompassed proliferating cells including those labeled by VZ/SVZ-associated enhancers MGE-*hs1538*, MGE-*hs1056*, and CGE- and LGE-*hs841*. The mitotic-associated enhancers captured both early VZ and SVZ cells. Cells in cl-1 expressed neural stem cell markers including higher expression of *Nfia*, *Nfib*, and *Olig2* (Figure 3D). Cl-3 was associated with higher levels of intermediate progenitor markers such as *Asb4* and *Ascl1*. Proliferative LGE and CGE cells from *hs841* comprised the majority of cl-2 and cl-9 and expressed VZ markers including *Lhx2* and *Fzd8*. Enhancer-negative LGE and LGE-*hs841* mitotic cells additionally form clusters cl-8 and cl-11, which expressed genes such as *Lrp4*, *Has1*, and *Kitl*. Cl-10 was composed primarily of enhancer-negative MGE cells and expressed *Lhx9* and *Tbr1*, indicative of cortical or diencephalic rather than basal ganglia identity. From random forest classification, markers that distinguished LGE, CGE, and MGE progenitor-associated enhancers recapitulated region-associated TFs from DC2 (Table S4). MGE rostrodorsal (*hs1538*) and caudoventral (*hs1056*) biased enhancers were distinguished by quantitative differences across TFs including *Otx2* and *Id4*, identifying TF expression gradients that distinguished progenitor cells across MGE regional axes.

Compared to proliferative enhancers, postmitotic enhancers active in SVZ/MZ mapped to distinct transcriptional clusters corresponding to emerging neuron types. LGE-*hs599* and CGE-*hs953* are both represented in cl-4, which expressed higher levels of genes including *Sp8*, *Tiam2*, and *Tshz1* (Figure 3D-E), suggesting cl-4 is more immature than cl-5 and cl-7 and has not yet acquired strong LGE or CGE regional specificity. Cl-5, composed predominantly of LGE-*hs599* cells, expressed genes including *Rbfox1*, *Tac1*, and *Zfp503*. Cl-7, composed predominantly of CGE-*hs953* and enhancer-negative cells, expressed genes such as *Six3*, *Tshz2*, and *Sema3a*. Genes defining these clusters shared general markers of early GABAergic projection neurons and were consistent with fate mapping of *hs599*^+^ and *hs953*^+^ cells to projection neuron populations in the adult forebrain, including striatal medium spiny neurons (McGregor et al., 2019; Silberberg et al., 2016) and Sp8^+^ neurons in the amygdala (Silberberg et al., 2016). MGE MZ-associated enhancers *hs192* and *hs799* also exhibited markers of early neuronal fate commitment (Figure 3D, Table S4). Cl-0, composed predominantly of *hs799*^+^ cells, expressed early MGE-derived cortical interneuron lineage markers including *Maf*, *Mafb*, *Nxph1*, *Calb1, and Sst*. Conversely, cl-6, composed of primarily *hs192*^+^ cells, expressed a wide range of TFs including *Tle4* and *Zic1, Zic3*, and *Zic4*, suggestive of GABAergic and cholinergic projection neuron commitment (Chen et al., 2010).

These experiments captured the cell-type specific activity of seven evolutionarily conserved enhancers across E11.5 GEs, showing the feasibility of enhancer-based genetic labeling paired with single cell transcriptomics and resolving lineage and spatial relationships via enhancer labeling. We found regionally separated but otherwise similar mitotic identities among enhancer-labeled progenitors across MGE, LGE, and CGE, and localized regional signatures within MGE via comparing dorsal (*hs1538^+^*) with more ventral *hs1056*^+^ cells. Across GEs, enhancer-labeled neuronal cell types emerged from more general early postmitotic clusters. Postmitotic cells in MGE separated into signatures suggesting GABAergic projection, cholinergic, and interneuron lineages differentially labeled by *hs192* and *hs799*. This initial survey defined specific cell populations and identified known and novel markers for maturation state and regional identity for progenitor and early born neuronal cell types, but had limited resolution to capture heterogeneity within enhancer-labeled cells or profile enhancer-labeled populations against all other cell types.

### Dissection of early born neuron types and positional identity in MGE via enhancer-labeled 3’ scRNA-seq and ISH

To more deeply interrogate emerging neuronal types labeled by *hs192* and *hs799* in E11.5 MGE, we performed scRNA-seq using the 10x Genomics Chromium system. Our initial scRNA-seq analysis of cells labeled by *hs192* and *hs799* suggested heterogeneity within these labeled early neuronal populations in the MGE. Indeed, fate-mapping experiments for these two enhancers (Silberberg et al., 2016) labeled other, MGE-derived GABAergic, cholinergic, and interneuron populations as well as early born CINs, populations that have not been the focus of previous scRNA-seq studies. FACS-purified MGE ungated and reporter-positive cells were dissected from embryos across three litters for each transgenic line and prepared for multiplexed scRNA-seq using MULTI-seq (McGinnis et al., 2019) (Figure 4A, Table S1). After quality control (Figure S5A-B), 4,001 single cells were used for downstream analysis. We used the same TF-curated approach developed for the C1 scRNA-seq dataset to drive cell clustering. Of the 463 TFs detected in 10x scRNA-seq MGE data, 421 (90.9%) were represented in both Fluidigm C1 and 10x Chromium systems (Figure 4B, S4F-I). We identified 18 cell clusters in the 10x dataset (Figure 4C). The largest determinant of transcriptional variation across cells was proliferative state (Figure 4D). Consistent with the C1 data, using TFs to drive cell clustering improved performance relative to highly variable gene approaches (Figure S5C-E). Overall, the 10x and MGE-derived C1 data had similar cluster topology and identified similar cell states, as determined by canonical correlation analysis of both datasets (Figure S6).

**Figure 4:**
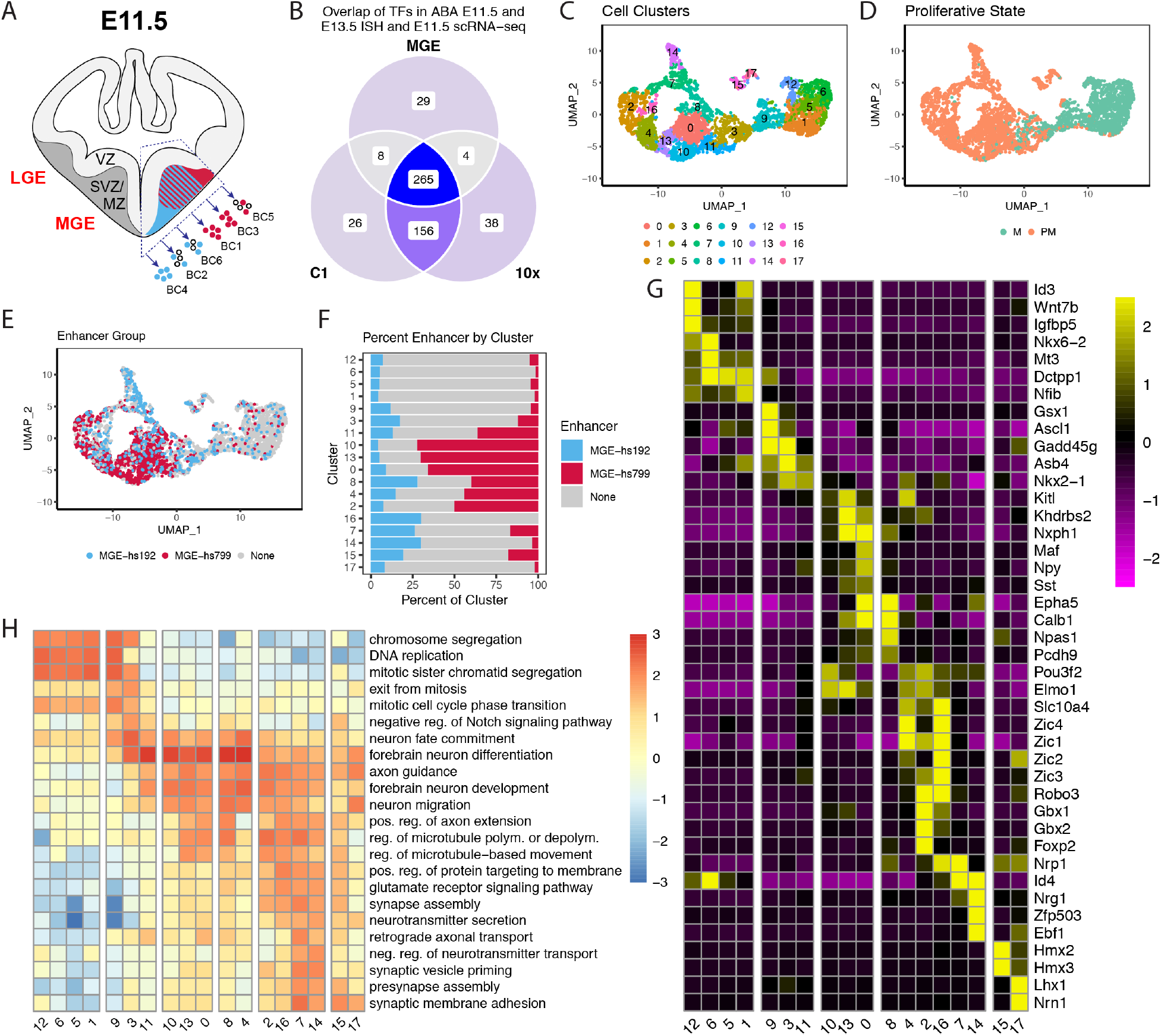
Regional and proliferative gradients within E11.5 MGE. (**A**) Schematic of dissections used for MULTI-seq (McGinnis et al., 2019), with colors corresponding to enhancer transgenic reporter expression in the medical ganglionic eminence (red: *hs799*; blue: *hs192*; open circles: unlabeled). LGE: lateral ganglionic eminence; MGE: medial ganglionic eminence; VZ: ventricular zone; SVZ: subventricular zone; BC: barcode. (**B**) Venn diagram depicting overlap of transcription factors scored within the Allen Developing Mouse Brain Atlas (ABA) for expression in E11.5 or E13.5 basal ganglia and transcription factors detected in 10x and C1 scRNA-seq datasets. (**C**) UMAP colored by TF-anchored cell clusters. (**D**) Visualization of single cells from E11.5 MGE by UMAP, colored by mitotic state (green: M, mitotic; orange: PM, postmitotic). (**E**) UMAP colored by enhancer (red: *hs799*; blue: *hs192*; grey: enhancer-negative). (**F**) Proportion representation of *hs192*, *hs799*, and enhancer-negative by cluster. (**G**) Representative differentially expressed genes by cluster. Color represents the mean Z-score of normalized expression across clusters. (**H**) Representative gene ontology biological process terms separating clusters. Color represents log2(observed/expected) for each term.

Cells were defined as enhancer-positive for *hs192* or *hs799* if they had at least one UMI (unique molecular identifier) count for CreER^T2^-IRES-GFP. Enhancer-positive cells from independent samples clustered together, demonstrating reproducibility across biological replicates and distinct distributions of enhancer-labeled *hs192* versus *hs799* cells (Figure 4E). The majority of enhancer-labeled cells mapped to post-mitotic clusters. *hs799*^+^ cells were substantially enriched (>50% of cluster composition) in cl-0, 10, and 13, and to a lesser extent in cl-2, 4, 8, and 11 (Figure 4F). In contrast, *hs192*^+^ cells were more broadly distributed, with the greatest representation in postmitotic clusters cl-7, 8, 14, 15, and 16, and decreased representation in cl-0, 2, 10, and 13 (Figure 4F). Cells labeled by at least one of the two enhancers were present in all postmitotic clusters alongside ungated enhancer negative cells.

We next performed differential gene expression (DE) analysis and Gene Ontology analysis across clusters using the full transcriptome of 18,088 genes to identify markers for each cluster and resolve maturation states and cell type identities (Figure 4G-H, Tables S3, S5). Proliferative clusters (cl-1, 5, 6, and 12), corresponding to cells within the MGE VZ, were enriched in terms including ‘DNA replication’ and ‘chromosome segregation’ (Figure 4H). In comparison, proliferative clusters cl-9 and cl-3, likely corresponding to cells within the SVZ, were enriched for terms such as ‘exit from mitosis’ (Figure 4H). Single cell resolution captured initiation of enhancer activity, which is first evident for both *hs799* and *hs192* among individual cells in late SVZ clusters based on transgene expression (Figures 4D-E). Postmitotic clusters subdivided into MGE-derived maturing GABAergic and cholinergic neuron and interneuron lineages (cl-0, 2, 4, 7, 8, 10, 11, 13, and 16), which are differentially labeled by *hs192* and *hs799*, and are described in detail below.

Some clusters were made up of postmitotic cells that appeared to originate outside the MGE (cl-14, 15, and 17), and were likely migrating through the MGE at E11.5 or captured at dissection boundaries. Cl-14 cells were enriched for *hs192*^+^ cells and had properties of LGE-derived immature medium spiny neurons, which express *Ebf1*, *Nrg1* and *Zfp503* (*Nolz1*), but not *Nkx2-1* (Figure 4G). Cl-15 contained both *hs192*^+^ and *hs799*^+^ cells and may derive from preoptic area (POA), based on *Hmx2* (*Nkx5-2*) and *Hmx3* (*Nkx5-1*) expression. Cl-17 cells were mostly transgene-negative, and may originate from regions adjacent to the subpallium, *e.g.* from the hypothalamus or prethalamic eminence. These non-MGE clusters are not further discussed.

### Projecting scRNA-seq cell identities onto developing mouse MGE via ISH data

Using transcriptional cell identities defined by scRNA-seq, we applied ABA ISH expression data (Lein et al., 2007) from the 689 TFs to map the anatomical distribution of cell populations in the E11.5 telencephalon. First, we manually graded expression of curated TFs in the VZ, SVZ, and MZ of MGE and/or LGE by reviewing individual sections from E11.5 and E13.5 in the ABA (Table S2). From this analysis we identified 332 genes with visually detectable mRNA ISH patterns in the BG, of which 283 were also detected in E11.5 scRNA-seq from C1 and 10x (Figure S4C). Following comprehensive review of genes with differential scRNA-seq expression and ABA ISH expression patterns in E11.5, E13.5, and E15.5 BG, we identified sentinel genes that could be used as spatiotemporal markers to presumptively assign cells or clusters to distinct neuroanatomical regions and/or specific neuronal lineages on the basis of their mRNA expression profiles. Using these sentinel markers and our scRNA-seq data, we characterized emergent cell lineages and regional distributions of proliferative (Figure 5) and postmitotic (Figure 6) cell populations in the MGE.

**Figure 5:**
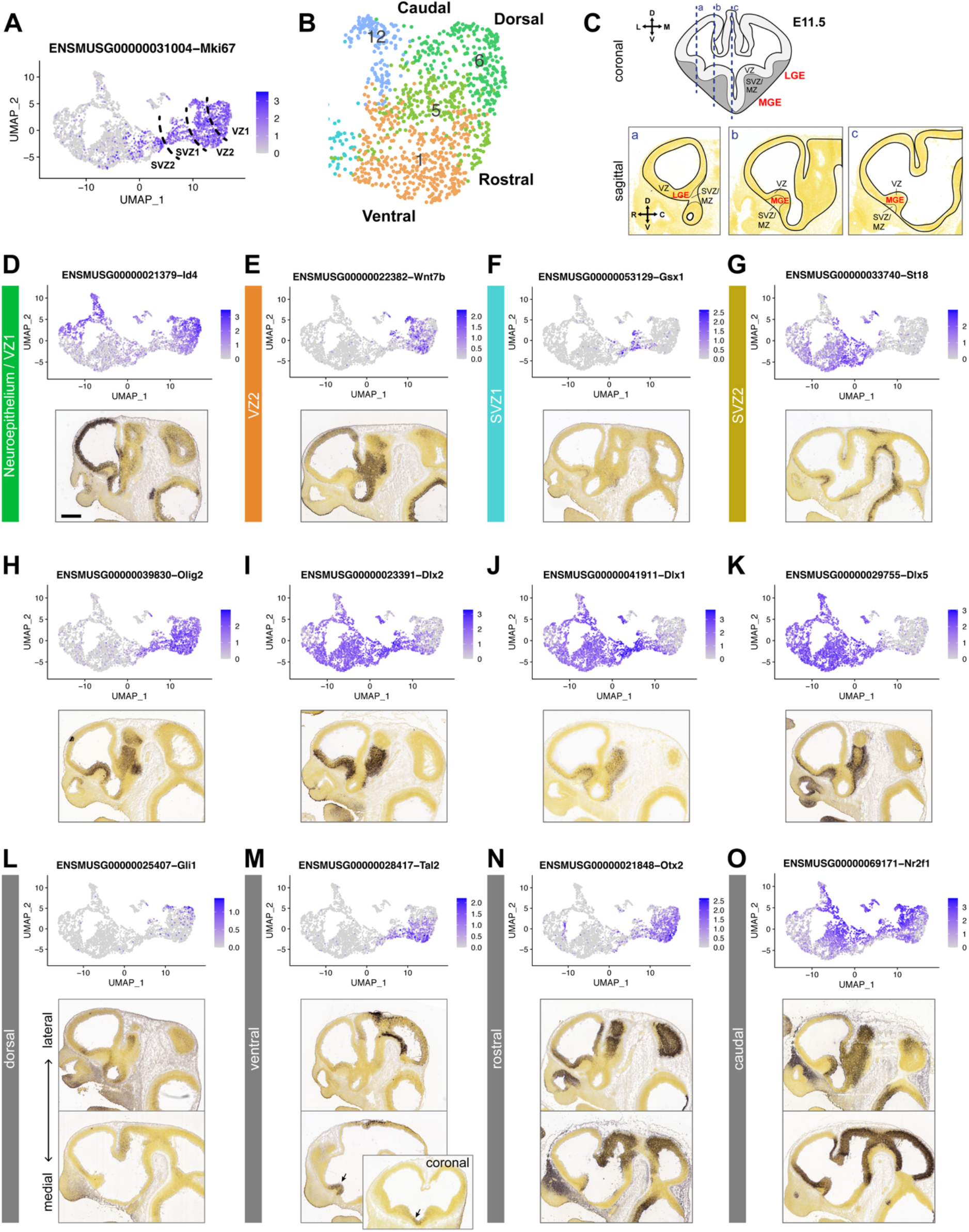
Separation of mitotic progenitors in E11.5 MGE. (**A**) Visualization of single cells from E11.5 MGE by UMAP, colored by expression of the mitotic marker *Mki67*. Dotted lines show proposed boundaries for cells from VZ1, VZ2, SVZ1 and SVZ2. (**B**) Region of UMAP covering the VZ cells, colored by TF-defined cell cluster to show proposed regional identities (also shown in Figure 4E). (**C**) Top: coronal schematic showing the positions (blue dotted lines) of parasagittal sections used to illustrate regional gene expression patterns. Bottom: reference images of parasagittal sections from the ABA with the relevant subpallial anatomical domains labeled. (**D-O**) Gene expression UMAPs and representative ISH on E11.5 medial parasagittal sections from the ABA showing expression domains for markers of various mitotic cell identities. (**D-G**) Gene markers for VZ-SVZ developmental stages (*Id4, Wnt7b, Gsx1, St18*). Colored labels correspond to main cell cluster identity for those cells. (**H-K**) UMAPS and ISH for *Olig2* and *Dlx2/1/5*, genes that help define SVZ1 and SVZ2 identity. (**L-O**) Gene markers with known regional expression patterns (*Gli1, Tal2, Otx2, Nr2f1*) in the MGE VZ correlate with spatially defined UMAP locations, corresponding to the schematic in (C). Later and medial sections are shown. (**M**) *Tal2* expression is highest at the medial level; coronal inset shows that this corresponds to the ventral MGE (arrow). Additional genes marking these developmental stages and regional identities are shown in Figures S7-8. Scale bar: 500 µm.

**Figure 6:**
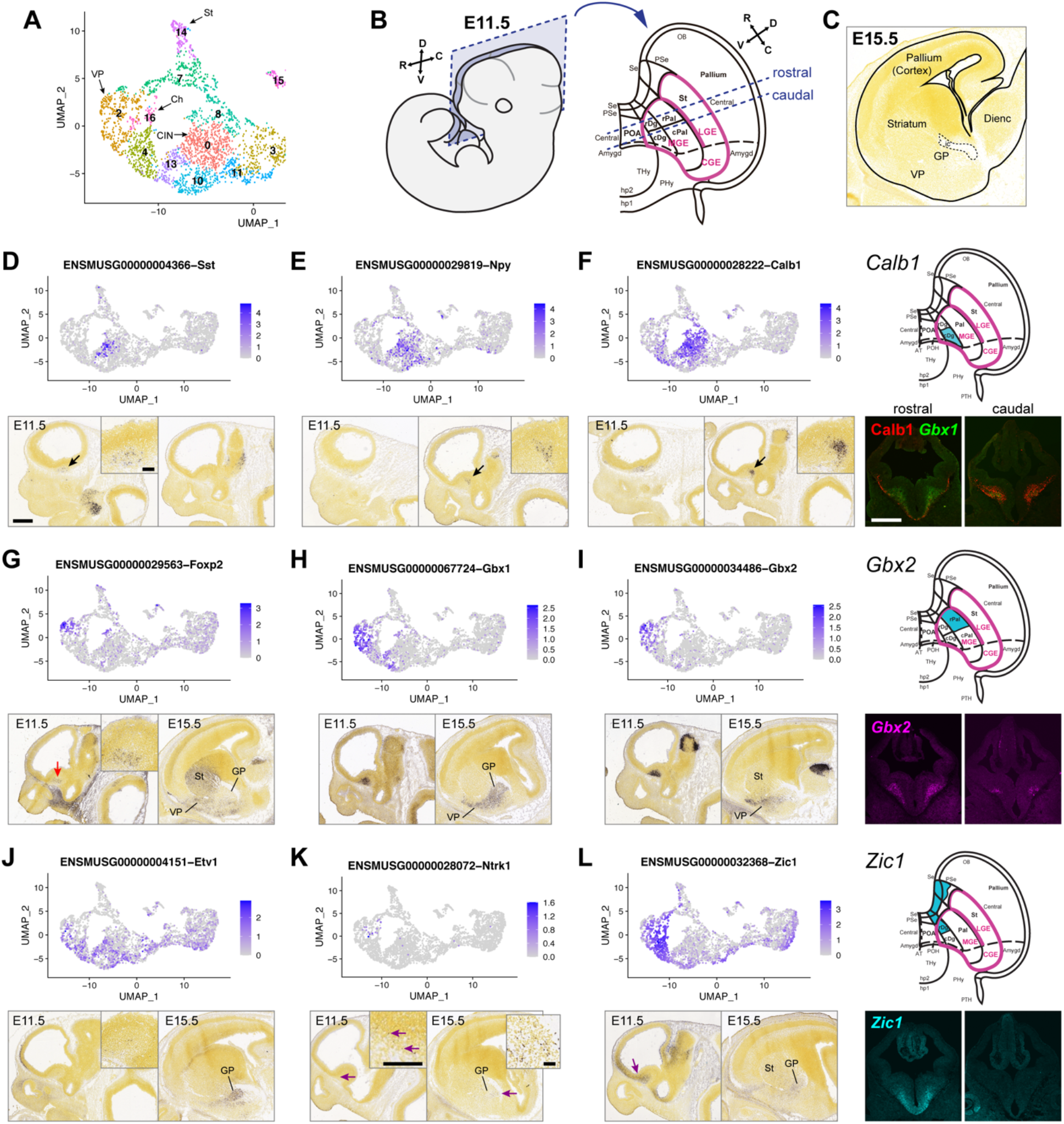
Separation of early neuronal lineages in E11.5 MGE. (**A**) Portion of UMAP colored by TF-defined cell cluster (also shown in Figure 4E) covering postmitotic cells, with proposed identities of cell clusters labeled. Abbreviations: VP, ventral pallidum; CIN, cortical interneurons; St, striatal medium spiny neurons; GP, globus pallidus; Amygd, amygdala; Ch, cholinergic. (B) Collapsed 2D topological map of the E11.5 MZ, as viewed from the sagittal section plane indicated. The schema is a variant of that in Silberberg et al., 2016. Note the addition of rostral and caudal Pal (Pallidum) and Dg (Diagonal area). See Table S6 for abbreviations. Blue dotted lines (rostral and caudal) indicate positions of coronal fluorescent images shown in **F**, **I** and **L**. (**C**) Schematic depicting broad anatomical domains at E15.5. VP: ventral pallidum; GP: globus pallidus; Dienc: diencephalon. (**D-L**) Gene expression UMAPs and representative ISH on E11.5 parasagittal sections from the ABA showing expression of markers for distinct postmitotic neuronal lineages. Arrows indicate regions of higher magnification insets and cells of interest. Arrow colors: black, CINs; red, VP; purple; Ch. For Calb1, Gbx2 and Zic1, a topological schematic illustrates the MZ expression domain for each gene, and coronal sections of fluorescent IHC (Calb1) or ISH (*Gbx1, Gbx2, Zic1*) are also shown to illustrate different MZ expression domains. Scale bars: low magnification, 500 µm; high magnification insets, 100 µm.

### MGE progenitors stratify by VZ to SVZ and dorsoventral and rostrocaudal axes

The analysis of mitotically active cells (*Mki67*^+^; Figure 5A) in the developing MGE provided evidence for distinct progenitor stages and regional patterning (Figure 5B-C). In our analysis, the assignment of four mitotic clusters was driven by genes previously associated with progenitor cell maturation steps, suggesting four discrete histogenetic stages: VZ1 (neuroepithelium), VZ2 (radial glial), SVZ1 (secondary progenitor 1), and SVZ2 (secondary progenitor 2) (Figure 5A). This interpretation is supported by mRNA ISH and scRNA-seq analyses, as described below.

The most immature VZ stage, the neuroepithelium (VZ1), was represented by cells with the highest expression of early mitotic markers such as *Hes1* and *Id4* (Kageyama et al., 2008) and roughly fit within cl-5 and cl-6 (Figures 5D, S7A). More mature VZ cells (VZ2; perhaps radial glia), organized as a diagonal zone in the UMAP plot, were characterized by expression of *Wnt7b, Id3*, and *Ttyh1;* they were roughly contained within cl-12 and the left part of cl-1 (Figures 5E, S7F,G). This zone also had high expression of *Hes5* and *Fabp7*, which mark radial glia (Feng et al., 1994; Kageyama et al., 2008) (Figure S7E,H). Other genes strongly marked both VZ zones (Eisenstat et al., 1999; Petryniak et al., 2007; Roychoudhury et al., 2020), including *Lhx2, Rest*, and *Rgcc* (Figure S7B-D). Genes like *Ascl1*, *Gsx2* and *Dlx2* began expression in the VZ as scattered cells both in the UMAP plot and by ISH (Figures 5I; S7J,K).

SVZ organized into the progressively more mature SVZ1 in cl-9 and SVZ2 in cl-3. We assigned SVZ1 identity to cl-9, based on overlapping *Olig2* and *Dlx2* expression (Petryniak et al., 2007) (Figure 5H,I). *Gsx1* showed perhaps the most specific SVZ1 periventricular expression by ISH and was largely confined to cl-9 (Figure 5F). Cells in this ‘isthmus’ cluster also expressed high levels of known SVZ genes (*Ascl1*, *Gsx2* and *Hes6*) (Long et al., 2009; Porteus et al., 1994; Roychoudhury et al., 2020) and neurogenic transition markers (*Btg2*) (Haubensak et al., 2004) (Figure S7J-L). *Gadd45g*, a marker of intermediate progenitors in the cortex (Yuzwa et al., 2017), was also highest in cl-9. SVZ2 identity was linked to cl-3. This most mature progenitor state was associated with the loss of *Olig2* and high levels of *Dlx1, 2, and 5* (Eisenstat et al., 1999; Petryniak et al., 2007) (Figure 5H-K). By ISH, cl-3/SVZ2 markers (e.g. *Prox1, Sp9* and *St18*) were expressed in a distinct layer of cells superficial to the SVZ1 markers (Figures 5G, S7M,N). *Insm1* and *Isl1* appeared to be expressed in both SVZ1 and SVZ2 (Figure S7O,P). Cl-8 and cl-11 may represent the earliest stage of neuronal commitment as SVZ2 cells exit the cell cycle; these clusters are discussed further below.

Leveraging canonical correlation analysis of MGE cells across both C1 and 10x datasets and ISH data, we found transcription factors and other markers that distinguished the regionally distinct *hs1538* and *hs1056* populations (Figure S6). Markers expressed in cells at the top of the UMAP plot (e.g. *Nkx6-2*, *Gli1* and *Gli2*) indicated dorsal MGE identity (Figures 5L, S8A,B). *Dach2*, a novel marker in this category, occupied a similar location in the UMAP plot and was expressed in the dMGE by ISH (Figure S8C). In the lower part of the UMAP plot, the genes expressed (*Shh, Slit2* and *Tal2*) indicated ventral (v)MGE identity (Hoch et al., 2015b) (Figures 5M, S8D-E). The highest expression of vMGE genes was observed in medial sagittal planes of ISH sections (Figure S8D-E). *Bcan*, *Dach1 and Sulf1* may also mark vMGE progenitor identity (Figure S8F-H). Molecular markers of MGE rostrocaudal position were also identified in the UMAP plot. The rostral MGE had high *Otx2* expression (Hoch et al., 2015b) whereas the caudal MGE was marked by high *Nr2f1* and *Nr2f2* expression (Hu et al., 2017) (Figures 5N,O, S8M). The preoptic area (POA) and pre-optic hypothalamus (POH) are contiguous with the caudal mitotic zone in the MGE. Cl-12 may represent a mixture of POA2 and POH progenitors, based on expression of *Nkx6-2, Dbx1*, and *Pax6* (Flames et al., 2007) (Figures S8B,O,P). POA1 cells lack these markers and have higher expression of *Etv1* (Flames et al., 2007); these cells may be intermixed with MGE cells within cl-1. The septum is contiguous with the rostral MGE, and the septal markers *Fgf15* (Borello et al., 2008), *Zic1 and Zic4* (Inoue et al., 2007; Rubin et al., 2010) were also expressed by rostral MGE progenitors (Figure S8I,J). *Pou3f1* and *Cntnap2* were novel markers of rostral cells (Figure S8K,L), whereas *Ptx3* was a new caudal MGE marker (Figure S8N). These patterning markers, including *Id4*, *Otx2*, and *Tcf7l2*, also distinguished *hs1538* (rostrodorsal biased) and *hs1056* (caudoventral biased) cells (Table S4), validating the regional identity evident among MGE progenitors. Thus, this enhancer labeling and TF-curated approach identified TF expression gradients capturing early MGE regional patterning among progenitors.

### Emergence of MGE neuronal lineages revealed by differential enhancer labeling

Postmitotic clusters were identified as the precursors of distinct MGE-derived cell lineages: GABAergic interneurons destined for the cortex (CINs) or other structures, GABAergic projection neurons, and cholinergic neurons (Figure 6A). Importantly, some of these cell types appeared to emerge in different spatial subdomains of the MGE based on marker gene expression. Early-born neurons emerge in SVZ2 and make up the MZ, which can be further visualized via a topological projection map which preserves spatial relationships across the telencephalon (Puelles et al., 2016; Silberberg et al., 2016) (Figure 6B-C). The organization and cellular outputs of the basal ganglia primordia identified herein using this topological map of E11.5 telencephalon correlate with progenitor zone anatomical map of (Flames et al., 2007) (Table S6). Previous scRNA-seq studies have captured some of these emergent neuronal identities but largely have not resolved the spatial organization underlying this process (Mayer et al., 2018; Mi et al., 2018). *Nkx2-1*, *Lhx6*, and *Lhx8* marked the MGE-derived postmitotic neurons (Figure 4G), and these cell groups were further resolved in UMAP plots and cluster assignments by their maturity state and presumed lineage. To analyze these relationships, we focused on major emergent interneuron, GABAergic projection neuron, and cholinergic lineages and the differential activity of *hs192* and *hs799* that distinguishes these specific emerging neuronal populations.

Cluster cl-0 and a subset of cl-13 were composed primarily of *hs799*^+^ and *hs192*^-^ cells and expressed immature interneuron markers. A subset of cl-8 included *hs799*^+^ and *hs192*^+^ that expressed *Mafb*. Cl-0 showed the most consistent expression of immature CIN markers including *Calb1*, *Cux2*, *Erbb4*, *Lmo3*, *Maf*, *Npy, Sox6, Sst and Zeb2* (Figures 4G, 6D-F, S9A-G). Earlier lineage markers such as *Lhx8* were reduced in these cells. Subsets of cl-13 and cl-8 cells were *Mafb*^+^, but lack *Maf* and other CIN markers. The small proportion of *Maf^+^ hs192*^+^ cells in cl-8 and cl-0 was consistent with our C1 data and in line with fate mapping of a subset of *hs192*-lineage cells to CINs (Silberberg et al., 2016). Sentinel genes for cells from cl-0 and cl-13 were expressed in a distinct area in the periventricular mantle zone of the caudal MGE based on ISH at E11.5. We found that *Calb1* expression is largely located in the caudal Diagonal (Dg) region, as illustrated on the topological map of the telencephalon (Figure 6F). The caudal Dg also showed expression of *Cux2*, *Erbb4*, *Npy*, *Sst*, *Maf*, and *Mafb*; the latter two genes are perhaps the most specific markers of MGE-derived CINs (Mckinsey et al., 2013; Pai et al., 2019) (Figure 6D,E, S9A-D). Contiguous with this region, in more lateral ISH sections, were cells expressing CIN markers, presumably migrating to the LGE and/or CGE on their way to the cortex (e.g. *Sst* and *Calb1* in Figure 6D,F). Additionally, *Adamts5*, *Bend4*, *Dlgap1, Kitl* and *Rai2* may be novel markers of immature CINs (Figures S9H,I). Based on putative maturation states and overlapping transcriptional signatures, immature interneurons from these three clusters can be presumptively mapped back to late progenitor cells in SVZ2 (cl-3) via cl-11 for cl-0, cl-10 to cl-11 for cl-13, and within cl-8. These findings map the neurogenic progression and distinct spatial niches for emerging early MGE-derived interneurons, including *hs799*^+^ cells that contribute to SST^+^ CINs, as well as other populations.

Of particular interest, the scRNA-seq signatures and corresponding ISH patterns suggested at least two distinct classes of GABAergic projection neurons differentially labeled by *hs799* and *hs192* and distinguished by *Gbx1^+^* or *Zic1*^+^ expression, respectively, that originate in different parts of the MGE (Figure 6H,L). The first GABAergic projection class was preferentially labeled by *hs799* and expressed *Gbx1*, with more restricted expression of *Foxp2* and *Gbx2*, and corresponded to *Shh*^+^ cl-13 cells and to subsets of cl-4 and cl-2 (Figures 6G-I, S8D). Based on a distribution on the UMAP, *Gbx1, Kitl, Lmo3, Sox6, Th, Tle4, Tshz2, Zeb2 and Zic1* (Flandin et al., 2010; Mckinsey et al., 2013) are additional markers for cells from cl-2, cl-4, and cl-13 (Figures 4G, 6L, 7D, S9E-H,J,K). These cells were presumptively mapped to less mature states in cl-10 and cl-1 (for cl-13 only) or cl-4 and cl-11 (for cl-2). On the topological map of the MGE, *Gbx2* expression was in the rostral portion of the pallidal (Pal) subdivision (Figure 6I). Anatomically, the E15.5 expression domains of *Foxp2*, *Gbx2*, *Gbx1 and Etv1*, determined by ISH, were nested along the pallidum’s radial axis. *Foxp2* expression was largely superficial, possibly in the ventral pallidum (Campbell et al., 2009) (Figure 6G). *Gbx1* expression encompassed the ventral pallidum and the entire globus pallidus (Figure 6H). *Gbx2* included the ventral pallidum and part of the globus pallidus (Chen et al., 2010) (Figure 6I). *Etv1* was expressed throughout the globus pallidus but in few cells in the ventral pallidum (Flandin et al., 2010) (Figure 6J). *Foxp2^+^* cells become superficial pallidal projection neurons (i.e. ventral pallidum), whereas other cl-2 cells, and some cl-4 and cl-13 cells, contribute to deeper pallidal structures (i.e. globus pallidus). Thus, we propose these populations that share increased *hs799* activity branch from immature neuron types within cl-2, cl-4, and cl-13.

**Figure 7:**
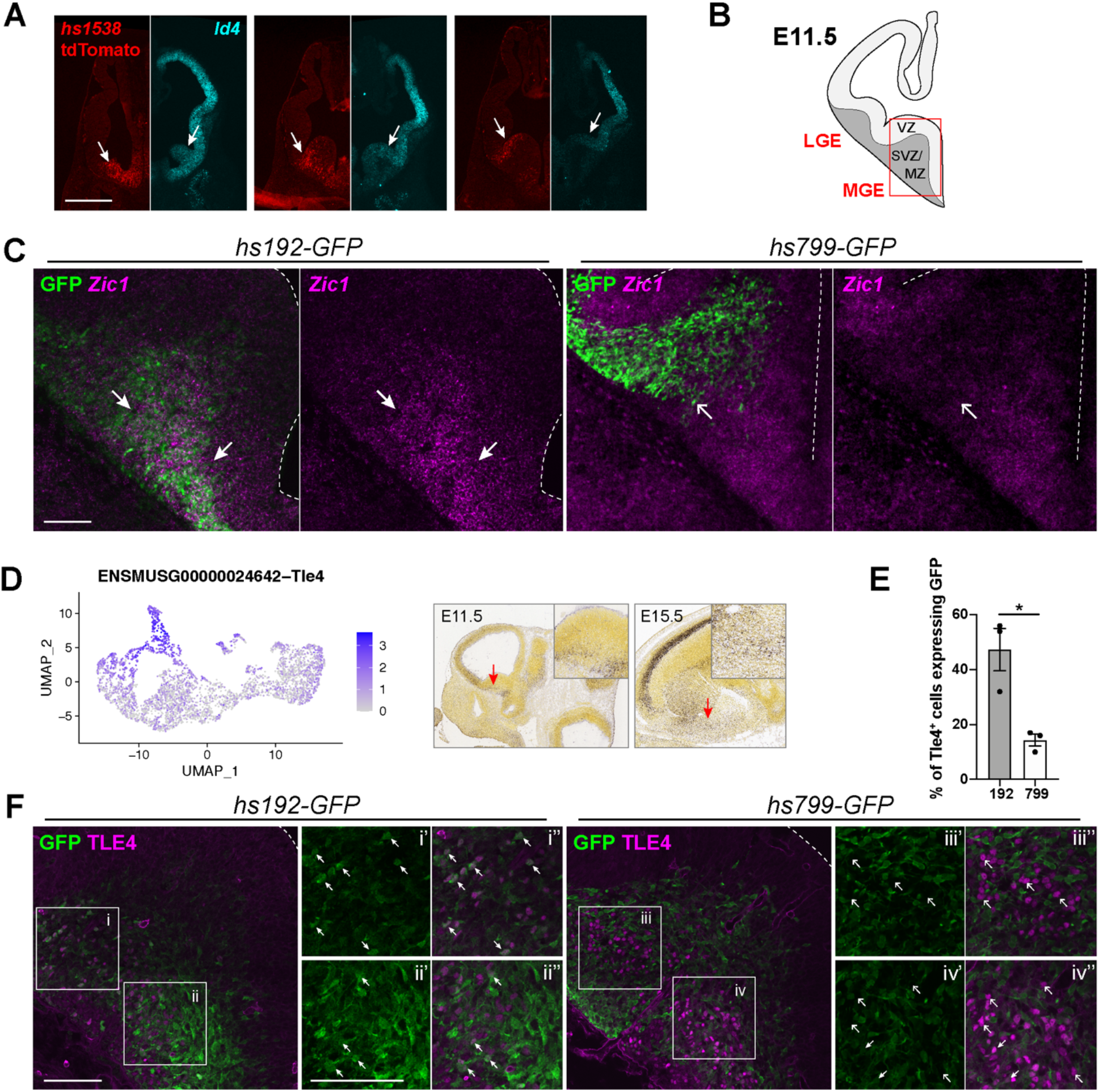
Spatial and epigenomic differences in E11.5 MGE enhancers. (**A**) Endogenous tdTomato expression (red) in *hs1538* E11.5 coronal sections after tamoxifen administration at E10.75, and *Id4* fluorescent ISH (cyan) on equivalent wildtype E11.5 sections. Arrows point to higher expression in the rostral and dorsal MGE VZ compared with ventral MGE. Scale bar: 500 µm. (**B**) Schematic showing location (red box) of low magnification images shown in (C) and (F). (**C**) *Zic1* fluorescent ISH (magenta) and GFP immunohistochemistry (green) on superimposed adjacent coronal sections of E11.5 *hs192* and *hs799* forebrain. Closed arrows indicate *hs192* GFP overlaps with *Zic1* expression in the ventral MGE. Open arrows indicate the ventral extent of *hs799* GFP cells, which does not extend into the region of high *Zic1* expression. (**D**) UMAP of Tle4 expression and representative ISH on E11.5 sagittal sections from the ABA. (**E**) Quantification of percent of Tle4 cells that are also GFP positive from (F). (**F**) Immunohistochemistry for Tle4 and GFP on *hs192* and *hs799* coronal sections. Double-positive cells are indicated by closed arrows and GFP-negative cells are indicated by open arrows. Scale bars: 100 µm. Ventricular surface of MGE is indicated by dotted lines.

The second GABAergic lineage expressed *Zic1*, as well as more restricted expression of *Zic3* and *Zic4*, and was enriched for cells with high *hs192* activity (Figure 6L). *Zic* TFs ISH MGE expression was restricted to the rostroventral MGE and overlapped little with *Gbx2*; *Zic1* may not overlap with the more caudal CIN markers (Figure 6D-F; S10A-D). On the topological map, *Zic1* expression was restricted to the rostral Dg region, and was continuous with *Zic1* expression in the septum (Figure 6L). As noted, *Zic1* was also expressed in the VZ and SVZ of the rostral MGE and septum, suggesting that this progenitor zone generates the *Zic1^+^* postmitotic cells in cl-11 and cl-4. It is unclear from ISH how *Zic1*-associated GABAergic projection neurons are spatially organized within the GP, though the spatial organization appears more diffuse across the GP. These *hs192*-biased *Zic1^+^* cells may constitute a distinct cell type within the GP.

Cells in cl-16 were exclusively enhancer *hs192*^+^ and *hs799*^-^ and expressed definitive cholinergic marker genes. Among these, *Ntrk1* (Sanchez-Ortiz et al., 2012) was the most specific to this cluster (Figure 6K). Although *Ntrk1* expression was weak in ISH at E11.5, scattered positive cells were visible in the striatum and GP by E15.5, consistent with the distribution of cholinergic interneurons (Figure 6K). Cl-16 cells also expressed *Gbx1, Gbx2*, *Isl1 and Zic4*, all of which are cholinergic lineage markers (Fragkouli et al., 2009; Chen et al., 2010; Magno et al., 2017; Asbreuk et al., 2002; Elshatory and Gan, 2008), as well as *Zic1, Zic2*, and *Zic5* (Figures 6H,I,L, S9L,M). *Fgf15* appeared to be a novel marker of this population, but its expression was not maintained as cells mature (Figure S9N). A subset of *hs192^+^* cl-4 cells also expressed some of these markers (e.g. *Zic2*) and may represent a less mature state or another type of cholinergic neuron. *Fgf8, Fgf17*, and *Nkx2-1/Zic4* fate mapping provides evidence that these cells arise from the junction of the rostromedioventral MGE with the septum (Magno et al., 2017; Hoch et al., 2015a), consistent with the expression of *Zic1*.

The final postmitotic group, cl-7, included two populations: the upper part was almost exclusively *hs192*^+^, while the lower part also contained hs799^+^ cells. The upper part of cl-7 was *Nkx2-1*^-^ and strongly expressed *Six3* and *Sp8* (Figure S9O,P), suggesting a CGE-derived CIN or LGE-derived olfactory bulb IN identity (Long et al., 2007). The lower part of cl-7 was *Nkx2-1*^+^ and *Lhx8^+^* (and thus likely MGE-derived); these cells also expressed genes shared by cl-2 and cl-14, including *Id4* and *Tle4* (Figures 5D, 7D). Earlier states of cl-7 cells appeared to map back to cl-8 then SVZ2 cl-3 cells, with *Nr2f2* expression in these clusters indicating caudal MGE origin (Figure S8M).

Across described postmitotic populations, enhancer-labeled scRNA-seq and ABA ISH indicated specific spatial localizations for cell types with divergent transcriptomic identities. To verify this finding, we performed fluorescent immunohistochemistry (IHC) and ISH to map protein expression across coronal sections for four sentinel genes representative of these populations (Figure 6F,I,L). We examined Calb1 for early CINs, predicted to be located in caudal Dg; *Gbx1* and *Gbx2* for GABAergic projection neurons in rostral Pal; and *Zic1* for rostral Dg GABAergic projection and cholinergic populations. These experiments validated that early born neuronal populations indeed exhibit spatial segregation to specific rostrocaudal and dorsoventral zones of the MGE MZ. Thus, our experiments captured novel lineage progression and associations between spatial and transcriptomic identity of E11.5 MGE-derived neuron populations.

### Validation of MGE enhancer-labeled progenitor and postmitotic populations

The combination of enhancer labeling and scRNA-seq paired with ISH defined distinct populations marked by progenitor (*hs1538* and *hs1056*) and postmitotic (*hs192* and *hs799*) enhancers in the MGE. We used ISH and co-labeling to validate transcripts that distinguish cells between these postmitotic and mitotic enhancer pairs. We found subtle but detectable transcriptomic differences between MGE progenitor cells labeled by *hs1538* and *hs1056* corresponding to regional spatial segregation of *hs1538* to dMGE VZ and *hs1056* to vMGE (Table S4). In contrast, there were broad transcriptional differences between the spatially intermixed populations of postmitotic cells labeled by *hs192* and *hs799* in the MGE (Tables S3,S4, Figures 3,5). Between *hs1538* and *hs1056*, among the most informative transcripts for differential enhancer labeling were *Id4*, *Tcf7l2*, *Zkscan1*, *Otx2*, and *Satb1* (Table S4). While the differential scRNA-seq signatures were subtle, ISH verified highest expression of the *hs1538*-associated marker *Id4* in the VZ of the dorsal and rostral MGE where *hs1538* is active (Figure 7A).

In postmitotic populations, *hs192* activity, which was biased toward GABAergic projection neurons in the striatum and cholinergic interneurons, was associated with higher levels of the transcription factor *Tle4* and several members of the *Zic* TF family including *Zic1*, *Zic3*, and *Zic4* (Table S4). Conversely, *hs799* activity favored populations of early CIN lineages expressing markers including *Mafb* and *Sst*, in addition to its activity in GABAergic projection neurons. As *hs192* and *hs799* MZ populations spatially intermingle, ISH alone was insufficient to verify specificity. Thus, to validate differential expression of genes across *hs192* and *hs799* labeled cells, we performed co-labeling of GFP^+^ enhancer-labeled cells and *Tle4* and *Zic1*, two transcription factors that were enriched in *hs192*^+^ cells (Figure 7B,C-F). As expected, enhancer-positive cells showed overlapping distributions in the MGE, but *hs192* activity and *Zic1* expression were highest in the rostroventral MGE and paraseptal region (Figure 7C,F). In addition, Tle4 protein was expressed in significantly more *hs192* GFP^+^ cells compared to *hs799* GFP^+^ cells (Figure 7E). Thus, ISH and co-labeling experiments validate the results from scRNA-seq-based separation of enhancer-labeled MGE cells.

## Discussion

Transcriptional profiling at single cell resolution has transformed our understanding of the diversity of cell types in the brain. While initial efforts to catalogue brain cell types have returned huge gains (Zeisel et al., 2018), new approaches are now needed to link transcriptional identity to location, function, and developmental lineage. Here, we combined novel enhancer-based cell labeling, TF-anchored clustering, and ISH-based spatial annotation to map the neurogenic landscape of embryonic mouse basal ganglia. Through this integrated approach, we illuminate enhancer activity in specific cells in vivo and provide new insights regarding the specification paths for early GABAergic neurogenesis in the ganglionic eminences.

Enhancer activity is often tightly restricted to specific cell types and developmental stages (Dunham et al., 2012; Nord, 2013; Reddington et al., 2020), making enhancers potent tools for genetic labeling and manipulation (Pattabiraman et al., 2014; Silberberg et al., 2016; Visel et al., 2013). However, the specificity and pattern of enhancer action in vivo at the single cell level has not been deeply explored. Most studies that examine enhancer specificity in the brain have used image-based assays of reporter expression (Visel et al., 2013) or orthogonal biochemical and epigenomic proxies to predict cell-type specific enhancer activity (Dunham et al., 2012). In contrast, our approach represents a major advance in resolution for in vivo function-based modeling of enhancer activity. Our results demonstrate that scRNA-seq can capture reporter transcripts across histogenetic subtypes labeled by individual enhancers, thus identifying enhancer-positive cells with high specificity even when enhancer activity was low or limited to specific cell types. scRNA-seq analysis revealed the onset and offset of enhancer activity as well as cell populations where each enhancer was active with fine-scale resolution. Overall, this study reinforces the specificity of activity across individual enhancers and increases resolution of enhancer activity mapping to offer a representative perspective of in vivo single cell enhancer activity for seven evolutionarily conserved neurodevelopmental enhancers.

Numerous genetic studies have probed mechanisms of BG development, with two recent studies (Mayer et al., 2018; Mi et al., 2018) using scRNA-seq to follow BG development and CIN lineage specification in E12.5-E14.5 embryonic mouse. Our results capture an earlier time when alternative neuronal lineages originate, using enhancer labeling, TF curation, and ISH to enable lineage tracking and spatial resolution of progenitor and postmitotic populations across the GEs. Similar to other studies, our results find transcriptomic differences among progenitor and postmitotic cells across the GEs, with additional markers of maturation and postmitotic lineages resolved via enhancer labeling. Further, integrating ISH data, we assigned likely identities to four types of MGE progenitors; distinct MGE progenitor regions; and enhancer-labeled subtypes of maturing MGE-derived interneurons, GABAergic projection neurons, and cholinergic neurons that together mature to form pallidal structures or migrate to become interneurons in the cortex, amygdala, and striatum.

Progenitor cohorts with overlapping transcriptional states and region-specific signatures were labeled by spatially distinct enhancers with VZ activity (*hs1538*, *hs1056*, *hs841*; Figures 2,3,4). Similar to published studies at older ages (Mayer et al., 2018; Mi et al., 2018), we found at E11.5 that scRNA-seq resolves the maturation gradient of BG progenitor populations from NSCs in the VZ to postmitotic neuronal precursors in SVZ2 (Figure 5A). Comparison of *hs1538^+^* and *hs1056^+^* enabled the discovery of genes whose expression in MGE progenitors defined rostrocaudal and dorsoventral axes. The rostrodorsal *hs1538*^+^ progenitors were enriched for *Id4*, *Otx2*, *Tcf7l2*, and *Zic1* (Figures 3, 5, 7; Table S4). In contrast, caudoventral progenitors were enriched for *Nr2f1/2* and *Tal2* (Figure 5). Interestingly, *Id4* is both associated with neuroepithelial progenitors (Bedford et al., 2005; Yun et al., 2004) and biased toward rostrodorsal MGE VZ, suggesting potential differences in progenitor state composition across regional domains within E11.5 MGE VZ. Of note, progenitors segregated on the UMAP across these axes when using a TF-anchored transcriptome analysis (Figure 5). We suggest that positional information in E11.5 MGE progenitors is largely encoded by gradients of these and other TFs rather than the expression of specific TF domains. Combinations of TFs then activate region-specific enhancers such as *hs1538*, as has been shown in classic model of ectoderm patterning in drosophila embryos (Levine, 2008) and in the cortical VZ (Pattabiraman et al., 2014).

Early-born CGE, LGE and MGE GABAergic projection, cholinergic, and inhibitory neuronal lineages are labeled by differential enhancer activities (*hs953, hs599, hs192, hs799*, respectively) (Figures 3,4,6). Compared to the distinct MGE signature, LGE and CGE cells had more similar transcriptomic identities, but diverged between *hs599* and *hs953* cells in later post-mitotic neuron clusters. Focusing on MGE, differential scRNA-seq comparison of enhancers *hs192* and *hs799* in combination with ISH annotation defined three spatially distinct regions in the MGE MZ that give rise to molecularly distinct cells. CINs (expressing multiple markers) were detected in the caudal Diagonal (Dg) region (Figure 6F). *Gbx1/2^+^* cells were mapped to the Pallidal (Pal) region and *Zic1^+^* cells to the rostral Dg regions (Figure 6I,L). We are intrigued by the possibility that *Gbx^+^* and *Zic^+^* cells contribute to distinct types of MGE GABAergic neurons, including within the GP, in addition to the cholinergic neurons already described (Chen et al., 2010; Magno et al., 2017). Previous studies have not identified a specific MGE region giving rise to CINs, nor spatially distinct zones for the generation of pallidal neurons and CINs. This contrast to our findings may be because our analysis focused on a younger age (E11.5) than most studies. Consistent with our results, (Puelles et al., 2016) previously showed that *Sst* expression begins in the Dg region. It is probable that as development proceeds, additional MGE regions also generate CINs, as suggested by many studies (Mayer et al., 2018; Mi et al., 2018; Silberberg et al., 2016). Overall, the results from these experiments define emerging GABAergic neuron types and elucidate relationships between spatial and transcriptomic characteristics of lineages in embryonic BG and in early mammalian brain development.

Beyond BG neurogenesis, our study has broader implications for the application of scRNA-seq to developing tissues. We demonstrate that anchoring scRNA-seq analysis on TF transcripts reduces the weight of cell cycle-driven and technical sources of variation (noise), improving the power of histogenetic cell type classification. Limiting analysis to TF transcripts greatly improved resolution of regional identity and maturation state for progenitors in the VZ and SVZ, consistent with the vast body of literature defining TF gradients as the master regulators of lineage specification in the BG and elsewhere. The extensive use of ISH annotations to define regional gradients enabled us to identify and interpret patterns of cell identity captured in scRNA-seq data. Finally, enhancer labeling enabled us to enrich, identify, and compare specific progenitor populations and early neuronal lineages, capturing novel transcriptional signatures for labeled lineages. Our results highlight the value and need for curated approaches in scRNA-seq analysis (e.g. our focus on TF transcripts) and the utility of leveraging orthogonal and accessory data, in this case ISH and enhancer labeling, to understand complex developmental processes. This study thus represents a new frontier, pairing scRNA-seq with functional analysis of enhancer activity in vivo and highlighting the utility of combining enhancer-mediated expression for labeling and characterization of specific cell types and offers insight into GABAergic neurogenesis in the embryonic mouse BG.

## ACKNOWLEDGEMENTS

Next-generation sequencing was performed at the Center for Advanced Technology at UC San Francisco and the UC Davis DNA cores. Confocal microscopy was performed at the UC San Francisco Nikon Imaging Center, supported by NIH S10 Shared Instrumentation grant 1S10OD017993-01A1. Cell sorting was performed at the UC San Francisco HDFCC Laboratory for Cell Analysis, using instrumentation supported by NIH grant P30CA082103. Additional cell sorting and C1 scRNA-seq were performed at the UCSF Parnassus Flow Core (RRID:SCR_018206) supported in part by Grant NIH P30 DK063720 and by the NIH S10 Instrumentation Grant S10 1S10OD021822-01. 10x scRNA-seq was performed by the UCSF IHG Genomics Core. L.S.-F. was supported by the UC Davis Floyd and Mary Schwall Fellowship in Medical Research, the UC Davis Emmy Werner and Stanley Jacobsen Fellowship, and by grant number T32-GM008799 from NIGMS-NIH. R.C.-P. was supported by a Science Without Borders Fellowship from CNPq (Brazil). This work was supported by the following research grants. J.LR.R.: NIMH R01 MH081880 and NIMH R37/R01 MH049428. A.S.N.: NIH/NIGMS R35 GM119831. Z.J.G.: Department of Defense Breast Cancer Research Program (W81XWH-10-1-1023 and W81XWH-13-1-0221), NIH (U01CA199315 and DP2 HD080351-01), NSF (MCB-1330864), and the UCSF Center for Cellular Construction (DBI-1548297), an NSF Science and Technology Center. Z.J.G. is a Chan-Zuckerberg BioHub Investigator. L.P. was supported by Seneca Foundation (5672 Fundación Séneca, Autonomous Community of Murcia) Excellency Research contract: 19904/GERM/15; project name: Genoarchitectonic Brain Development and Applications to Neurodegenerative Diseases and Cancer.

## AUTHOR CONTRIBUTIONS

L.S.-F. and A.N.R. are listed as joint first authors, as each led components of the experiments and analysis. J.L.R.R. and A.S.N. are listed as joint senior and corresponding authors. L.S.-F., A.N.R., S.N.S., J.L.R.R., and A.S.N. designed the experiments. Dissections, single-cell preparations and histology: A.N.R. and S.N.S.; scRNA-seq library preparation: L.S.-F., A.N.R., and I.Z.; bioinformatics: L.S.-F., R.C.P., K.J.L., C.S.M., T.E.R., Jr., and A.S.N.; in situ hybridization prioritization and scoring: S.N.S., G.L.M., M.H., C.T., and H.Z. Topological map: L.P. and J.L.R.R. L.S.F., A.N.R., J.L.R.R., and A.S.N. drafted the manuscript. All authors contributed to manuscript revisions.

## Methods

### RESOURCE AVAILABILITY

#### Lead Contact

Further information and requests for resources and reagents should be directed to and will be fulfilled by the Lead Contacts, Alex S. Nord and John L. R. Rubenstein.

#### Materials Availability

The enhancer transgenic mouse lines used in this study have been previously published (Silberberg et al., 2016) and deposited to the MMRRC repository.

#### Data and Code Availability

The datasets generated during this study are available on GEO (accession TBD). The analysis codes used for this study can be found on the Nord Lab Git Repository (https://github.com/NordNeurogenomicsLab/).

### EXPERIMENTAL MODEL AND SUBJECT DETAILS

#### Mice

The enhancer transgenic mouse lines used in this study have been previously published (Silberberg et al., 2016) and deposited to the MMRRC repository. All animal care, procedures, and experiments were conducted in accordance with the NIH guidelines and approved by the University of California, San Francisco animal care committee’s regulations (Protocol AN180174-02). Pregnant dams were housed in mating pairs, or singly housed with additional environmental enrichment. Mice were housed in a temperature-controlled environment (22-24°C), had ad libitum access to food and water, and were reared in normal lighting conditions (12-h light-dark cycle). Embryos of either sex were used, and all embryos of the correct genotype from a single litter were pooled as a single biological replicate for all sequencing experiments.

We used mice from 7 previously published enhancer transgenic lines (Silberberg et al., 2016): *hs192-CreER^T2^-IRES-GFP, hs599-CreER^T2^-IRES-GFP, hs799-CreER^T2^-IRES-GFP, hs841-CreER^T2^-IRES-GFP, hs953-CreER^T2^-IRES-GFP, hs1056-CreER^T2^-IRES-GFP* and *hs1538-CreER^T2^-IRES-GFP*, herein referred to simply by their enhancer ID, *e.g.*, *hs192*. Enhancer line hemizygous transgenic male mice were mated to CD-1 wildtype or *Ai14* tdTomato Cre-reporter female mice (MGI ID: 3809524) (Madisen et al., 2010) to obtain embryos for experiments. All transgenic mice were maintained on a mixed background outcrossed to CD-1. For inducible tdTomato labeling of enhancer positive cells with the *hs1538* line, *Ai14* reporter female mice were mated to enhancer transgenic males and dosed with 55 mg/kg tamoxifen dissolved in corn oil at 6 pm the day before embryo harvest (E10.75 for harvest at E11.5). 14 mg/kg progesterone was included to improve embryo survival.

### METHODS DETAILS

#### C1 scRNA-seq

##### Cell isolation

Pregnant dams were sacrificed by CO_2_ inhalation, confirmed by cervical dislocation. E11.5 embryos were removed and placed into ice-cold Earle’s Balanced saline solution (EBSS). Transgene-positive embryos were identified by screening on a fluorescent microscope. In a clean dish with ice-cold EBSS, the MGEs, LGEs or CGEs were dissected out and placed into a 1.5 mL Eppendorf tube containing EBSS on ice. Tissue was pooled from all transgene-positive embryos in a single litter.

The tissue was dissociated in 300 µL of 0.25% trypsin-EDTA solution supplemented with 10 U/mL recombinant DNase I (Roche) for 15 minutes at 37°C. Trypsinization was stopped by addition of 300 µL DMEM with 10% FBS, and the tissue was gently triturated 10-15 times with a P1000 pipette and filtered through a 40 µm filter to achieve a single-cell suspension. The cells were spun down for 3 minutes at 500 rcf and resuspended in FACS buffer (EBSS + 0.5% BSA + 2 mM EDTA). DAPI (50 ng/mL final concentration) was included to stain dead cells.

For gated samples, GFP- and tdTomato-positive cells were isolated by fluorescence activated cell sorting on a FACSAria II flow cytometer (BD Biosciences). FACS gating was set using a transgene-negative sample, and DAPI-positive dead cells were excluded. Single-cell sorting mode was used to maximize sample purity. For ungated samples, DAPI-stained cells were excluded but GFP-positive and GFP-negative cells were collected.

##### C1 cell capture and cDNA generation

For unsorted and ungated experiments, cells were counted on a haemocytometer and diluted to 150-300 cells/µL in FACS buffer. The cell mix was prepared and loaded onto a Fluidigm integrated fluidics chip (C1™ Single-Cell mRNA Seq IFC, 5–10 µm, #100-5759) on the Fluidigm C1 system according to the manufacturer’s instructions.

For GFP- and tdTomato-gated samples, cells were sorted directly into the loading well of the IFC (integrated fluidic circuit) according to the manufacturer’s note (https://www.fluidigm.com/articles/cell-sorting-directly-to-the-c1-ifc). Briefly, the IFC plate was placed on the plate chiller of the FACSAria sorter, the inlets were covered with PCR-plate sealing film, and the sort stream was directed to the cell loading well. The film over the cell loading well was removed and 3 µL of C1 cell suspension buffer was added to the well. 1,500 cells were then sorted directly into the well. Finally, 1.5 µL of FACS buffer was added to give a final volume of 7.5 µl and a concentration of 200 cells/µL. The cell suspension was pipetted gently 2-3 times to mix, and cells were loaded onto the IFC using the C1 machine according to the manufacturer’s instructions. After capture, each well of the IFC was visually examined on a Keyence microscope and scored for the number of cells and presence or absence of cell debris.

Cell lysis, reverse transcription and cDNA amplification were performed on the Fluidigm C1 machine according to the manufacturer’s mRNA-seq protocol using the SMARTer Ultra Low RNA Kit for the Fluidigm C1 System (Takara Bio #634833). cDNA amplicons (∼3 µL) were harvested into a 96-well plate containing 10 µL C1 DNA dilution buffer per well, as described in the protocol, and stored at −80°C for library preparation.

##### C1 scRNA-seq library preparation

1 uL of diluted cDNA per well was used for library preparation using the Nextera XT DNA Sample Prep Kit (Illumina, #FC-131-1096). Each C1 IFC was pooled into one library, for a total of up to 96 samples per library. Sequencing library quality was assessed using the high-sensitivity dsDNA assay in an Agilent Bioanalyzer.

#### Multiplexed 10x scRNA-seq library generation (MULTI-seq)

E11.5 MGE tissue was dissected as described above and dissociated using the Papain Dissociation System (Worthington) with a modified protocol. MGEs pooled from a single litter were incubated with 200 µL papain solution supplemented with 10 U/mL recombinant DNase I (Roche) for 10 minutes at 37 °C on a rocking platform, then spun down for 3 minutes at 300 rcf. The papain solution was replaced with 200 µL ice-cold EBSS, and the tissue was gently triturated ∼10 times with a P1000 pipette to achieve a single-cell suspension. The cells were spun down and resuspended in 200 µL EBSS for fluorescence activated cell sorting on a FACSAria II machine (BD Biosciences). An aliquot of cells was stained with trypan blue and counted on a haemocytometer. The cell suspension was diluted if necessary with EBSS, aiming for ≤500,000 cells in a volume of 200 µL.

Dissociated cells were labeled with barcoded lipid-modified oligonucleotides (LMOs) as previously published (McGinnis et al., 2019), using a different barcode for each single-litter pooled MGE sample. Excess LMOs and papain were quenched by adding 1 mL of ovomucoid/BSA inhibitor (Worthington). Barcoded cells were spun down for 5 minutes at 500 rcf and resuspended in 300 µL EBSS with 1% BSA. DAPI (50 ng/mL final concentration) was included to stain dead cells. Just prior to sorting, cells were passed through a 40 µm filter to remove any remaining clumps.

GFP-positive cells were isolated by FACS as described above. Cells were sorted into Lo-Bind Eppendorf tubes containing EBSS with 1% BSA. Sorted, barcoded cell samples were then pooled, spun down, and resuspended in a small volume of EBSS with 1% BSA ready for processing on the Chromium 10x system.

Single-cell cDNA libraries were generated using the Chromium Single Cell 3ʹ GEM, Library & Gel Bead Kit (v3, PN-1000075) according to the manufacturer’s instructions. After the first cDNA clean-up step with 0.6x SPRI beads, the supernatant containing the barcode library fraction was saved and processed as described previously (McGinnis et al., 2019). The barcode library (5%) and cDNA library (95%) were pooled for sequencing on an Illumina NovaSeq SP lane.

#### C1 sequencing, alignment, and gene expression quantification

Libraries for the C1 scRNA-seq samples were sequenced on an Illumina HiSeq 4000 instrument using a single-end 50-bp protocol. Reads were uniquely aligned to the mouse genome (GRCm38, modified to append a custom chromosome containing the individual sequences of the transgenes CreER^T2^, IRES, EGFP, and tdTomato) using STAR (v2.7.0e) (Dobin, 2013), and read duplicates were removed using the Picard tools function MarkDuplicates (v2.18.4) (*Picard Toolkit, 2019*). Gene counts were generated using subread featureCounts (v1.6.3) (Liao et al., 2014), to ENSEMBL GRCm38 release 95, using a customized gtf annotation file containing annotations for the four transgenes. Gene counts were normalized to gene length and library size using reads per kilobase of transcript, per million mapped reads (RPKM) or to counts per million mapped reads (CPM) for downstream analysis and visualization.

#### 10x sequencing, alignment, and gene expression quantification

Libraries for 10x MULTI-seq samples were sequenced on an Illumina NovaSeq SP instrument. The raw 10x data was processed using CellRanger (v3.0.2) (Zheng et al., 2017) using the custom mouse genome described above. Samples were demultiplexed using the MULTI-seq (McGinnis et al., 2019) pipeline (https://github.com/chris-mcginnis-ucsf/MULTI-seq). Briefly, raw FASTQ files were split into cell barcode, UMI, and sample barcode sequences, then reads were parsed to generate a sample barcode UMI count matrix. The cell barcode matrix was put through the MULTI-seq classification workflow to identify single cells from unclassified and doublet/multiplet cells. A second round of semi-supervised classification was performed to reclassify negative cells for potential false negatives. The cell by gene UMI count matrix was loaded into Seurat (v3.2.2) (Butler et al., 2018; Stuart et al., 2019) as a Seurat object, and normalized using the Seurat function *SCTransform* (Hafemeister and Satija, 2019) with default parameters for both full transcriptome and TF-curated analyses.

#### C1 quality control

Raw read quality was assessed using FastQC (v0.11.7) (Andrews, 2010). Aligned library quality was assessed using RSeQC (v2.6.4) (Wang et al., 2012) for 5’ or 3’ bias, exonic read distribution and GC content distribution. We also assessed quality of individual samples by comparing the following distributions: total uniquely aligned reads, total assigned reads, total expressed genes, percent mitochondrial reads, percent ribosomal genes, and percent pseudogenes. Samples with greater or less than 2 standard deviations from the mean in any of the above metrics were discarded. In addition, samples scored as ‘two or more cells’ during visual inspection were removed from analysis. Pseudogenes, ribosomal genes (ribosomal subunit and rRNAs), mitochondrial genes, and six sex-linked genes (*Xist*, *Eif2s3x*, *Kdm6a*, *Ddx3y*, *Eif2s3y*, *Kdm5d*) (Armoskus et al., 2014) were removed from the data matrix.

We used the R package Seurat (v3.2.2) (Butler et al., 2018; Stuart et al., 2019) for feature selection, clustering and visualization. We compared two normalization approaches for quality control and downstream analysis. RPKM-normalized data was used to create two Seurat objects, with minimum.cells set to 10, and minimum.genes set to 2,000. We applied a manual log1p transformation to the RPKM data to generate normalized data in the ‘data’ slot of the Seurat object. We applied regression for FACS-processing state and total mapped reads on the gene count data using *ScaleData*. To define proliferative state and cell cycle phase, we used the Seurat function *CellCycleScoring* on a published list of cell cycle genes (Kowalczyk et al., 2015) and calculated G1/S and G2/M phase scores for each cell. We then calculated the difference between G1/S and G2/M to define whether a cell was mitotic (M) or post-mitotic (PM).

We also compared our results to those generated with raw count data normalized using the NormalizeData function with scale.factor set to 1e6 to generate CPM-normalized data. Results were largely similar between RPKM- and CPM-normalized data and only the RPKM-normalized results are reported. We note that short genes were more likely to be considered highly variable using CPM-normalized data (data not shown).

#### 10x quality control

We used the MULTI-seq pipeline described above to remove negative or multiplet cells from downstream analysis. After subsetting data to only singlets, we used the full transcriptome data to perform cell clustering with Seurat, which split cells into two populations based on mitochondria RNA (mtRNA) expression: one population with high mtRNA expression (>5% total UMIs) and one with low expression (<5% total UMIs). We removed all cells from the high mtRNA expression population, and repeated cell clustering using the cleaned cells. We identified one outlier cluster likely corresponding to hematopoietic cell lineages based on expression of hemoglobin genes, and removed that cluster, resulting in 4,001 cells for downstream analysis.

#### C1 transcription factor (TF)-based clustering

From the full dataset, we extracted counts for 689 transcription factors with mRNA in situ hybridization data available in the Allen Developing Mouse Brain Atlas (Lein et al., 2007) that were scored for expression in E11.5 and E13.5 basal ganglia and cortex (Table S2). Of the 689 scored transcription factors, 455 were expressed in >10 cells in our dataset. These 455 genes were used for dimensionality reduction using principal components analysis (PCA) and Uniform Manifold Approximation and Projection (UMAP) (McInnes et al., 2018) as visualization tools. Following PCA, we used jackstraw analysis with 100 iterations to define statistically significant (*p-value* < 0.05) principal components (PCs) driving variation. The first 13 significant PCs were used to define clusters using *FindNeighbors* (using k.param = 5, nn.method = “annoy”, annoy.metric = “euclidean”) and *FindClusters* (using resolution = 1.2, algorithm = 2, group.singletons = F). To generate the UMAP visualization we used Seurat’s *RunUMAP* (reduction = “pca”, n.neighbors = 15, n.epochs = 1000, negative.sample.rate = 10). Using this approach, we defined 12 clusters.

#### 10x transcription factor (TF)-based clustering

We used a similar approach as described above for the C1 clustering. Of the 689 scored transcription factors, 463 were expressed in greater than 10 cells. These 463 genes were used for downstream processing. The first 14 significant PCs were used to define clusters using *FindNeighbors* (using k.param = 15, nn.method = “rann”, annoy.metric = “euclidean”) and *FindClusters* (using resolution = 1.4, algorithm = 2, group.singletons = F). To generate the UMAP visualization we used Seurat’s *RunUMAP* (reduction = “pca”, n.neighbors = 20, n.epochs = 500, negative.sample.rate = 15). Using this approach, we defined 18 clusters.

#### Differential expression analysis

For differential expression (DE) analysis across TF-based clusters in both C1 and 10x datasets, we expanded to all expressed genes and used Seurat’s *FindAllMarkers*. Briefly, one cluster of cells was compared to all other cells using a Wilcoxon rank-sum test. We only considered genes that were expressed in at least 25% of cells in either population and had log_e_-fold-change greater than 0.25 between populations. Genes with adjusted *p*-values < 0.05 were considered statistically significant. We also performed DE analysis between enhancer groups (e.g. MGE-*hs192* vs MGE-*hs799*, or MGE-*hs192*-PM vs MGE-*hs799*-PM) using the same parameters.

#### Diffusion mapping pseudotime analysis

We performed diffusion mapping via the R package *destiny* (Angerer et al., 2016) on the TF-based C1 dataset. Briefly, the log-normalized TF-curated RPKM data was passed to the destiny function *dm*, which generates 50 diffusion components and corresponding cell eigenvalues. We assessed differential expression along the diffusion components using a generalized additive model via the *gam* function in R.

#### Random forest classification

We used random forest (Breiman, 2001) via the R package *randomForest* on the TF-based C1 dataset to identify transcription factors important for accurately classifying cells in pairwise comparisons of enhancers. Briefly, we ran *randomForest* independently 10 times using the log-normalized TF-curated RPKM C1 data matrix, using 5,000 trees per forest, for each of the pairwise comparisons (MGE-*hs1056*-M vs dMGE-*hs1538*-M; MGE-*hs192*-PM vs MGE-*hs799*-PM; CGE-*hs841*-M vs LGE-*hs841*-M; and CGE-*hs953*-PM vs LGE-*hs599*-PM). We extracted the out-of-box error rates for all forests in addition to the mean decrease in accuracy (MDA) and mean decrease in node impurity (MDNI) scores for all genes, and calculated mean and standard deviation. Genes were ranked by the mean MDA and MDNI scores.

#### Gene ontology analysis

We used the R package topGO (v2.36.0) (Alexa and Rahnenfuhrer, 2020) to perform gene ontology (GO) analysis by TF-based cluster and enhancer group on DE genes in the 10x dataset. Mouse GO data were downloaded from Bioconductor (org.Mm.eg.db) (Carlson, 2019). We restricted analysis to GO Biological Process annotations and required a minimal node size of greater than 20. We used the ‘weight01’ framework and Fisher test to define significant terms (*p-value* < 0.05). The background for all enrichment comparisons is all 18,088 expressed genes. For Figure 5, clustering of selected significant GO terms was performed using the R package pheatmap (Kolde, 2015) using *hclust* and default parameters.

#### Full transcriptome analysis

We repeated clustering analysis of single cells using the full transcriptome for both C1 and 10x scRNA-seq datasets. To define highly variable genes (HVGs), we calculated the mean of all expressed genes in the Seurat object data slot using the Seurat function *FindVariableGenes* using default parameters. The top 3,000 HVGs were used for dimensionality reduction using PCA and UMAP visualization. For C1 data, we used the first 10 PCs to define 11 clusters using *FindNeighbors* (using k.param = 15, nn.method = “rann”, annoy.metric = “euclidean”) and *FindClusters* (using resolution = 1, algorithm = 2, group.singletons = F). For 10x data, we used the first 17 PCs to define 17 clusters using *FindNeighbors* (using k.param = 20, nn.method = “rann”, annoy.metric = “euclidean”) and *FindClusters* (using resolution = 1, algorithm = 2, group.singletons = F). UMAP visualization was generated using *RunUMAP* with the same parameters described for each dataset above. DE analysis was performed across clusters as described above.

#### Combined analysis of C1 and 10x datasets

We used canonical correlation analysis (CCA) to combine the C1 and 10x datasets using both TF-curated and full transcriptome datasets. Briefly, the raw counts for both datasets were merged together, excluding genes that exhibited expression only in one dataset, and normalized using *NormalizeData* (normalization.method = “LogNormalize”, scale.factor = 1e6). Integration anchors between both datasets were identified using *FindIntegrationAnchors* (reduction = “cca”, dims = 1:50, normalization.method = “LogNormalize”, anchor.features = 3000). Datasets were integrated using 50 CCA components from *FindIntegrationAnchors* with *IntegrateData* (dims = 1:50), and center-scaled data generated using *ScaleData*. The integrated dataset was used for principal components analysis and the first 15 PCs were used to define 17 clusters using *FindNeighbors* (using k.param = 5, nn.method = “rann”, annoy.metric = “euclidean”) and *FindClusters* (using resolution = 1, algorithm = 2, group.singletons = F). UMAP visualization was generated using *RunUMAP* (reduction = “pca”, n.neighbors = 20, n.epochs = 1000, negative.sample.rate = 10).

#### ABA ISH transcription factor scoring

Expression levels of all transcription factors in the Allen Developing Mouse Brain Atlas (Lein et al., 2007) were estimated in subdomains of the subpallium for two ages: E11.5 and E13.5. The subpallium was divided into 4 anatomical domains (LGE, MGE, Septum, and POA). The LGE and MGE were further subdivided into laminar subdomains: VZ/SVZ and MZ for E11.5; VZ, SVZ1, SVZ2, and MZ for E13.5. Expression was annotated from sagittal sections in each subdomain as an Intensity value and a Density value on a 0-5 scale. The intensity value was derived from the color-coded intensity viewing option called “expression mask” on the ABA. The density value was assigned by binning the frequency of individual signals across subdomains onto the 0-5 scale.

#### Histology

##### Section preparation

Pregnant dams were sacrificed by CO_2_ inhalation, confirmed by cervical dislocation. E11.5 embryos were removed and placed into ice-cold 1x PBS. The heads were cut off and drop-fixed in 4% paraformaldehyde in PBS overnight at 4 °C. The fixed heads were then cryoprotected in 20% sucrose in PBS overnight at 4 °C, embedded in OCT compound (TissueTek) and frozen on dry ice. Cryostat sections 15 µm thick were cut directly onto SuperFrost slides and allowed to dry.

##### Fluorescent immunohistochemistry

Sections were rinsed in PBS, blocked for 1 hour at room temperature in PBST (PBS + 0.25% Triton X-100) with 10% FBS, and incubated with primary antibody diluted in PBST + 10% FBS overnight at 4 °C. Primary antibodies used were mouse anti-Tle4 (1:100, Santa Cruz Biotechnology, #sc-365406, RRID: AB_10841582), rat anti-GFP (1:1000, Nacalai Tesque, #04404-84, RRID: AB_10013361) and rabbit anti-GFP (1:5000, Abcam, #ab6556, RRID: AB_305564). The following day, sections were washed 2 x 15 min in PBST and 2 x 15 min in PBS, incubated for 2 hours at room temperature with Alexa fluor-conjugated secondary antibodies (1:750, Invitrogen). Finally, sections were washed 2 x 15 min in PBST and 2 x 15 min in PBS and coverslipped with Fluorescence Mounting Medium (DAKO #S3023).

##### Fluorescence in situ hybridization (FISH)

The antisense RNA probes used in this study have been described previously (Long et al., 2009). In situ hybridization was performed on 15 µm cryostat sections as described previously (Lindtner et al., 2019) up until the antibody blocking step, with the addition of a peroxidase quenching step for 20 minutes in 3% H_2_O_2_ in PBS after the SSC washes. After blocking, slides were incubated with anti-Digoxigenin-POD Fab fragments (Roche #11207733910) diluted 1:500 in NTT blocking buffer for 1 hour at room temperature. Slides were then washed for 3 x 5 minutes in NTT and developed with Cy5-Tyramide signal amplification reagent (TSA Plus Kit, Akoya Biosciences #NEL745001KT) according to the manufacturer’s instructions. Finally, sections were washed 3 x 15 minutes in PBS, incubated for 5 minutes with nuclear counterstain Hoechst 33342 (1:1000 in PBS, ThermoFisher #H3570), rinsed 3 x 5 minutes in PBS, and coverslipped with Fluorescence Mounting Medium (DAKO #S3023).

##### Imaging

Low-magnification epifluorescent images were taken using a Coolsnap camera (Photometrics) mounted on a Nikon Eclipse 80i microscope using NIS Elements acquisition software (Nikon) and a 4x or 10x objective. Confocal images were taken with 20x air and 40x oil objectives on an Andor Borealis CSU-W1 spinning disk confocal mounted on a Nikon Ti Microscope and captured with an Andor Zyla sCMOS camera and Micro-Manager software (Open Imaging). The raw images were pre-processed with ImageJ software (v2.0.0) to adjust brightness/contrast and convert to 8-bit RGB. Confocal images were stitched laterally to create composites using the Grid/Collection stitching ImageJ plugin with linear blending (Preibisch et al., 2009).

##### Cell counts

Cell counting was performed on single confocal image planes in ImageJ using the Cell Counter plugin. Counts were summed from at least 3 rostrocaudal sections for each brain. Tle4-positive cells were counted first with the GFP channel hidden, excluding cells in the VZ (designated by nuclear staining), then scored as positive or negative for overlap with GFP staining.

## Supplementary Figures

**Supplementary Figure 1:**
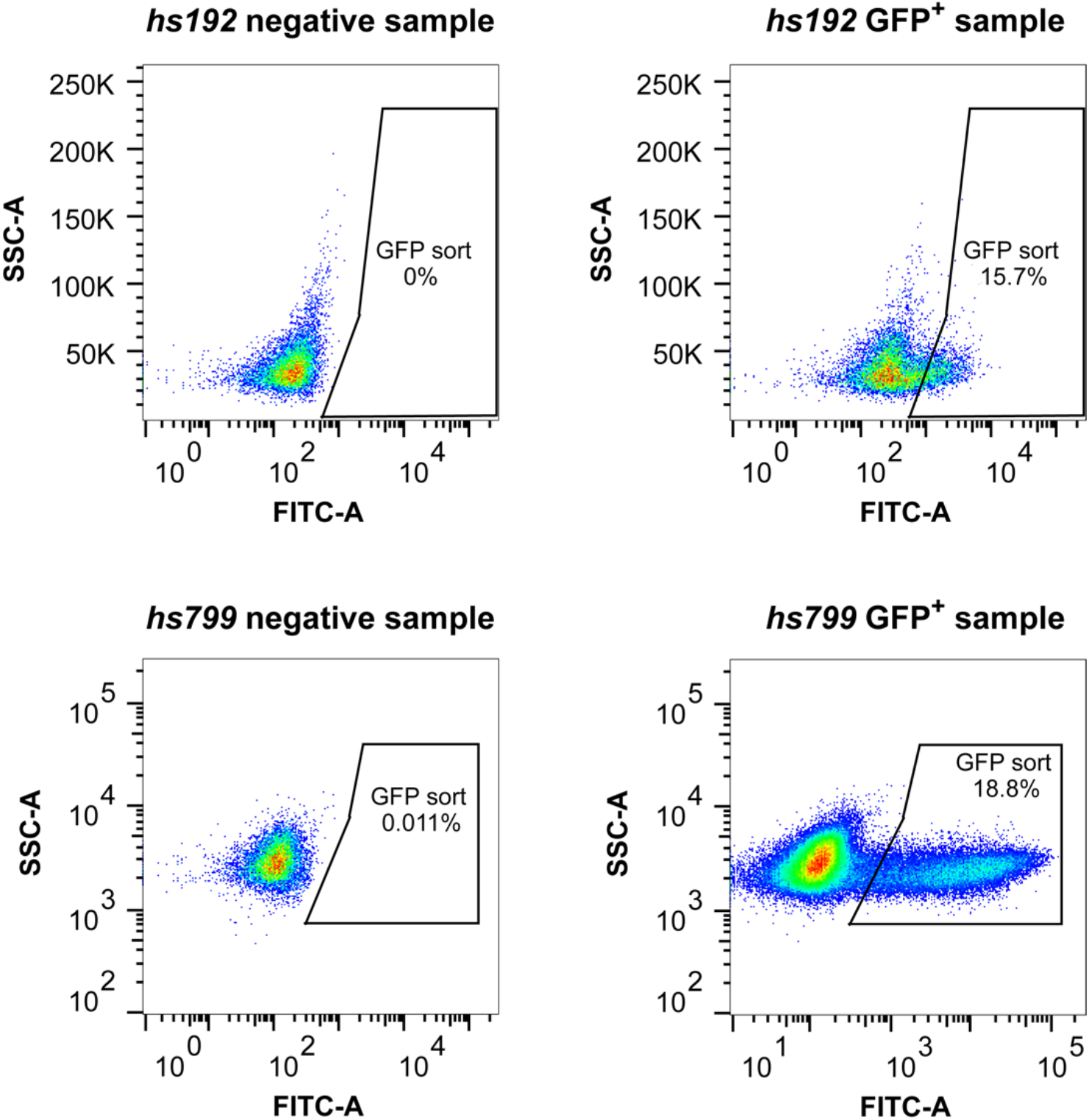
FACS gating schematics. Density plots showing side-scatter (SSC) versus GFP fluorescence intensity (FITC) for negative and GFP-positive samples from *hs192* and *hs799* MGE. GFP sort area shows cells collected for scRNA-seq and the percentage of live cells captured. Negative samples were used to set the gating. *hs799*^+^ cells show brighter GFP fluorescence than *hs192^+^* cells.

**Supplemental Figure 2:**
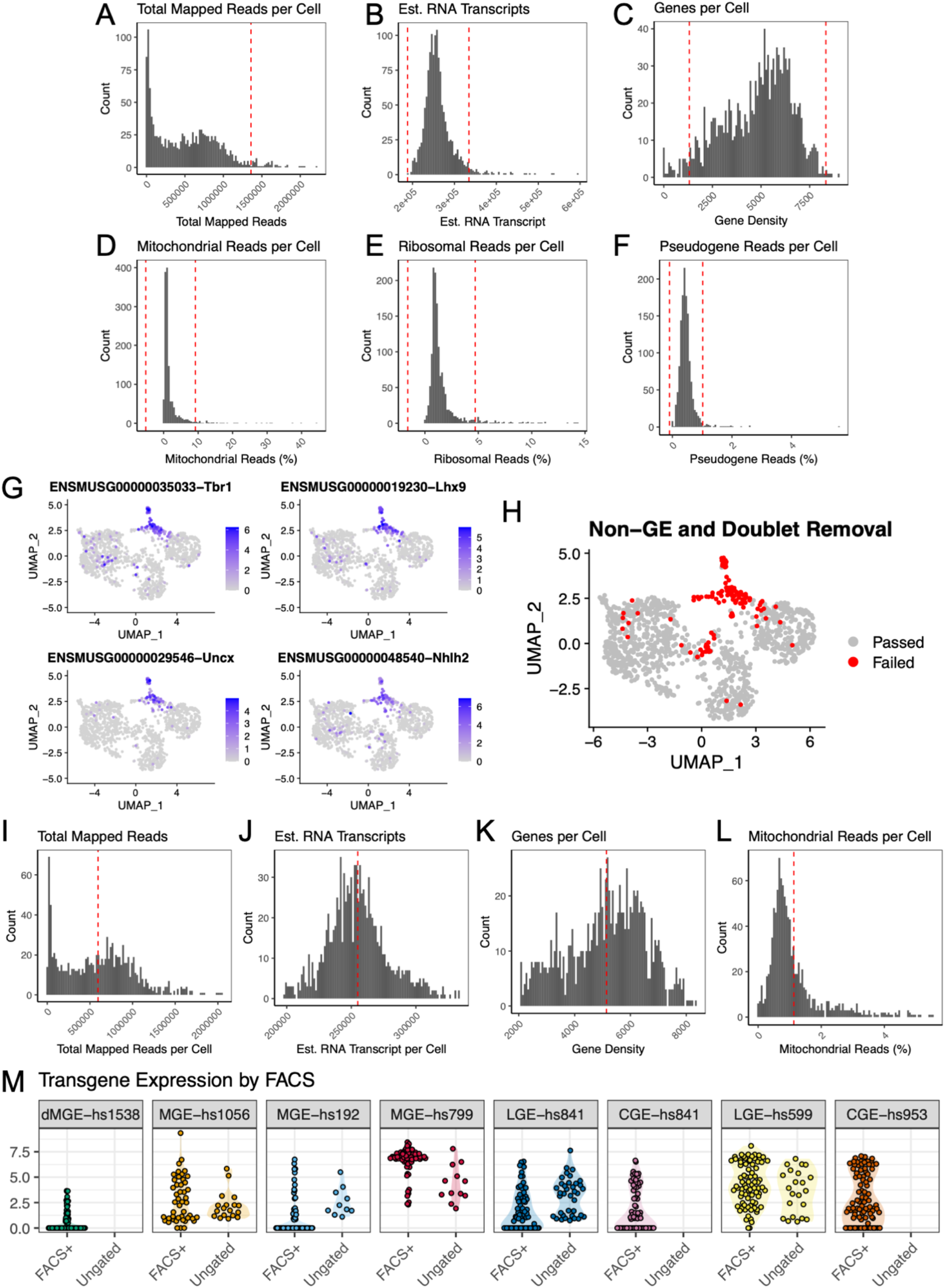
C1 scRNA-seq quality control. (**A-F**) Histogram of total uniquely mapped reads (**A**), estimated RNA transcript (**B**), number of expressed genes (**C**), percent mitochondrial reads (**D**), percent ribosomal reads (**E**), and percent pseudogenes (**F**) per cell, post-duplication removal for all sequenced samples. Cells above or below the red lines were removed from downstream analysis. Red lines represent mean ± 2 s.d. (**G**) UMAP of transcription-factor-curated clusters after removal of cells flagged in A-F, colored by normalized expression of four genes used for flagging cells as non-ganglionic in origin (*Tbr1*, *Lhx9*, *Uncx*, and *Nhlh2*). (**H**) UMAP colored by quality control flag for contaminating cells and suspected doublets (passed: used for downstream analysis; failed: removed from downstream analysis). (**I-L**) Histograms of total mapped reads (**I**), estimated RNA transcripts (**J**), expressed genes per cell (**K**), and percent mitochondrial reads per cell (**L**), for all cells that passed final quality control. Red line represents mean. (**M**) Normalized transgene expression for each of the seven enhancers profiled, separated by FACS^+^ gating or no FACS^+^ gating (if performed).

**Supplemental Figure 3:**
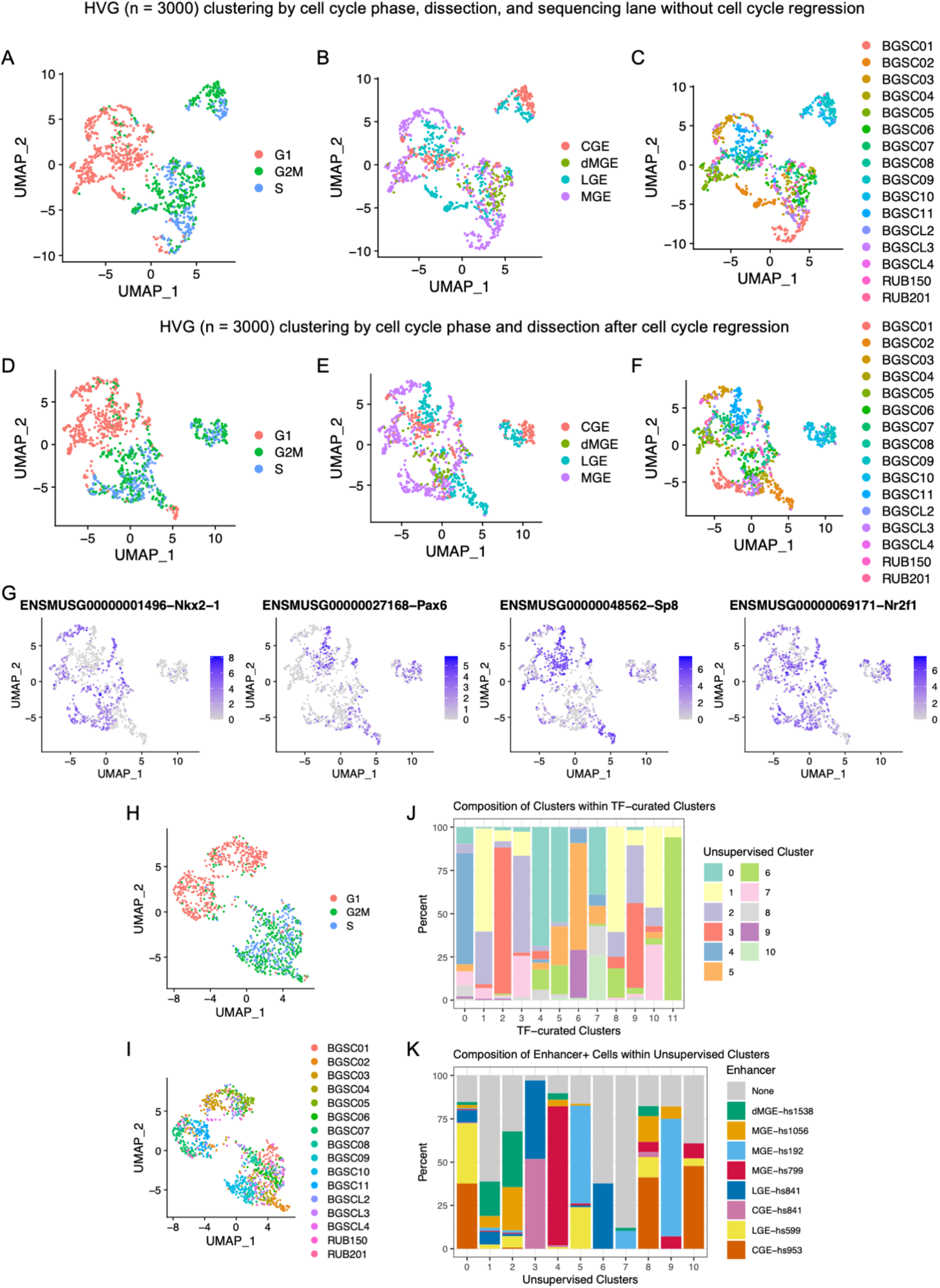
Comparison between full transcriptome and TF-curated analyses for C1 scRNA-seq. (**A-C**) UMAP representations using full transcriptome (“unsupervised”) clustering of the top 3,000 highly variable genes (HVGs) with no cell cycle phase regression, colored by cell cycle phase (**A**), region of dissection (**B**), and sequencing lane (**C**). (**D-F**) UMAP representations using full transcriptome of the top 3,000 highly variable genes (HVGs) after cell cycle phase regression, colored by cell cycle phase (**D**), region of dissection (**E**), and sequencing lane (**F**). (**G**) UMAP colored by normalized expression of four region-defining transcription factors (*Nkx2-1*, *Pax6*, *Sp8*, and *Nr2f1*). (**H**) TF-curated UMAP colored by cell cycle phase. (**I**) TF-curated UMAP colored by sequencing lane. (**J**) Bar plot of TF-curated clusters by percent representation from unsupervised (full transcriptome) clusters. Some unsupervised clusters (e.g. cl-10, light green) remain consistent between clustering methods, while others (e.g. cl-1, light yellow) are split across multiple TF-curated clusters. (**K**) Bar plot of unsupervised clusters by enhancer representation. Some enhancers (e.g. MGE-*hs799*, red) are relatively consistent across clustering methods, while others (e.g. MGE-*hs192*, light blue) split across clusters.

**Supplemental Figure 4:**
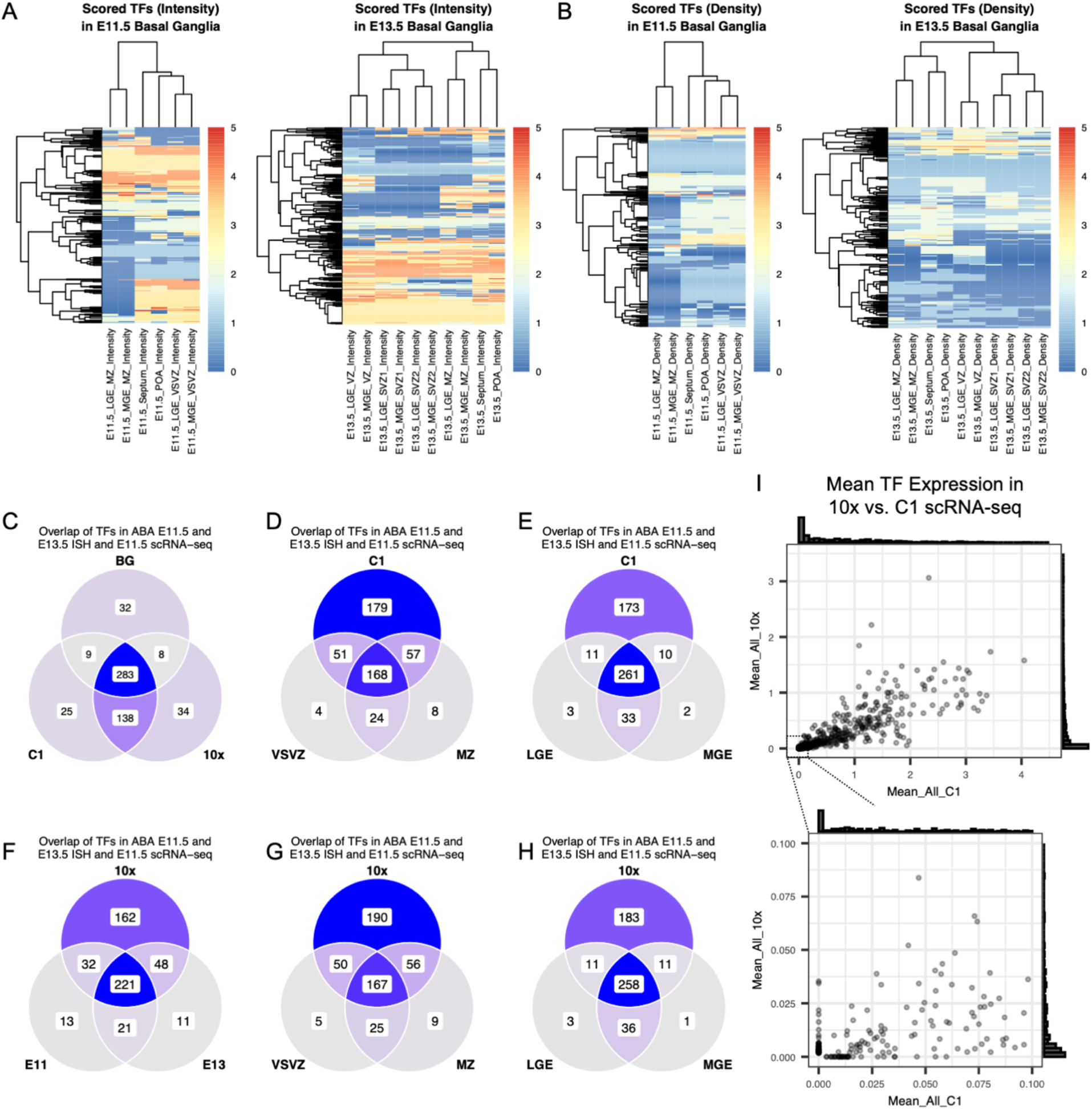
Allen Developing Mouse Brain Atlas and scRNA-seq overlaps. (**A**) Heatmap for E11.5 (*left*) and E13.5 (*right*) scored transcription factor (TF) expression by intensity in Allen Developing Mouse Brain Atlas (ABA) RNA in situ hybridization (ISH) for 689 TFs. Rows: transcription factors; columns: areas and time points scored. MGE: medial ganglionic eminence; LGE: lateral ganglionic eminence; POA: preoptic area; VSVZ: ventricular/subventricular zone; MZ: mantle zone. (**B**) Heatmap of scored expression as in (A), by density. (**C**) Venn diagram of overlap of TFs with non-zero intensity score in E11.5 and E13.5 basal ganglia (MGE and LGE combined) from the ABA dataset (“BG”) to detected TFs in C1 and 10x scRNA-seq datasets. (**D-E**) Overlap of TFs expressed in C1 scRNA-seq, separated by non-zero VSVZ vs MZ intensity scores (**D**) or LGE vs MGE scores (**E**) in E11.5 and E13.5 ABA ISH. (**F**) Overlap of TFs expressed in 10x scRNA-seq, separated by non-zero intensity scores in E11.5 vs E13.5 ABA ISH. (**G-H**) Overlap of TFs expressed in 10x scRNA-seq, separated by non-zero VSVZ vs MZ intensity scores (**G**) or LGE vs MGE scores (**H**) in E11.5 and E13.5 ABA ISH. (**I**) (*Top*) Plot comparing mean normalized expression of transcription factors between 10x and C1 scRNA-seq. Cell distribution along x- and y-axes are represented by histograms. (*Bottom*) Enlargement of the bottom-left corner of the top panel, showing distribution of low-representation TFs and dataset-specific expression.

**Supplemental Figure 5:**
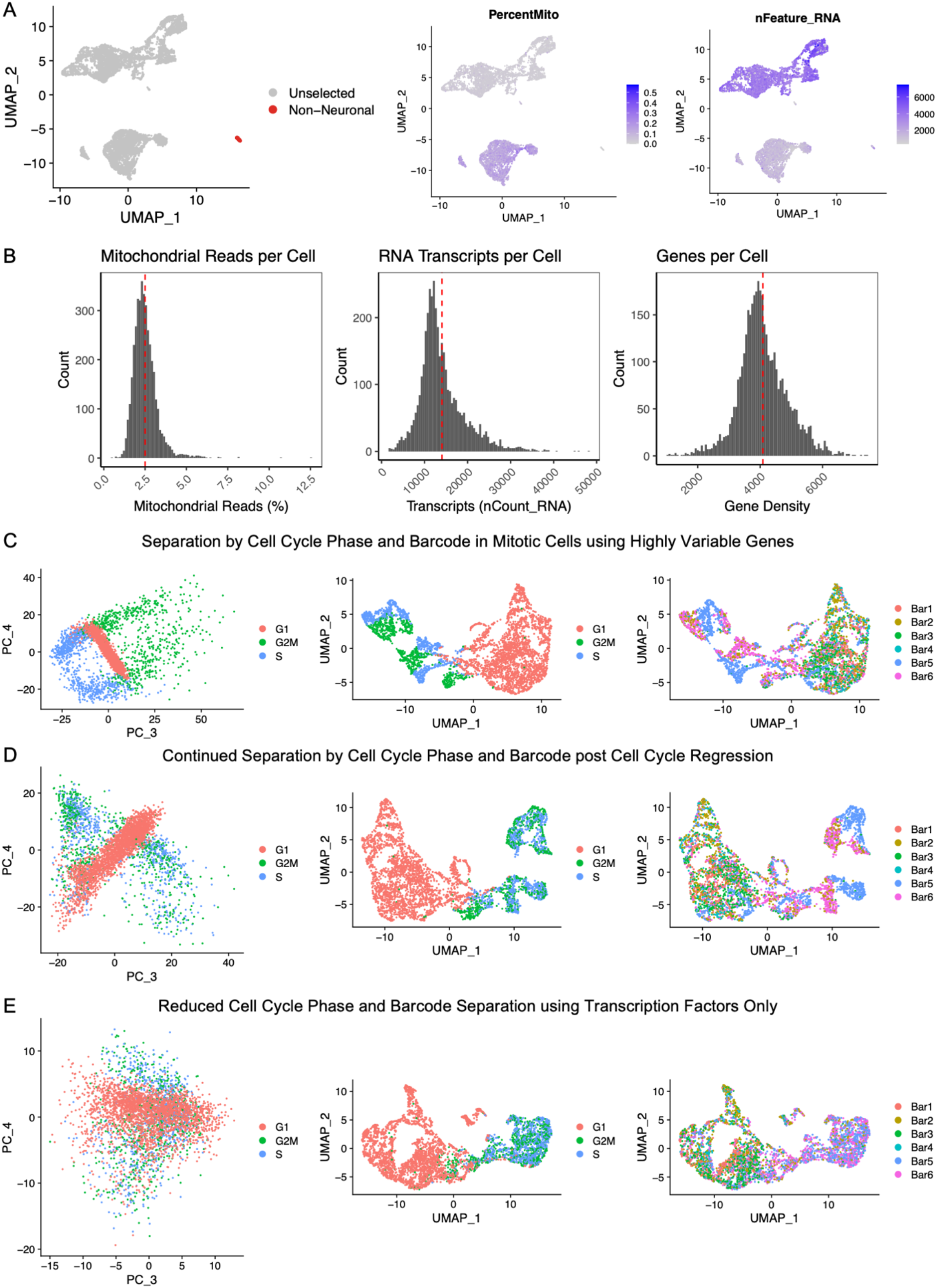
10x scRNA-seq quality control. (**A**) UMAP representation of single cells after sample demultiplexing and removal of multiplets using the MULTI-seq pipeline, colored by cells flagged for likely non-neuronal origin (*left*), percent mitochondrial reads, (*middle*), and number of expressed genes per cell (*right*). (**B**) Distribution of percent mitochondrial reads (*left*), estimated RNA transcripts (*middle*), and number of expressed genes per cell (*right*) with means (red line) after removal of all cells flagged by quality control metrics (effectively cells UMAP_2 < 0 in A). (**C**) Principal components (PCs) 3 and 4 separate mitotic cells by cell cycle phase in full transcriptome (unsupervised) clustering using the top 3,000 highly variable genes (HVGs) (*left*). In UMAP representation of unsupervised clustering, cells separate by cell cycle phase (*middle*) and MULTI-seq barcode (*right*) in mitotic cells. (**D**) After cell cycle regression, cell cycle phase separation is negated (*left* and *middle*) but cells still separate by barcode in mitotic cells (*right*). (E) Using transcription factor-based clustering, both cell cycle phase separation (*left* and *middle*) and barcode separation (*right*) are negated.

**Supplemental Figure 6:**
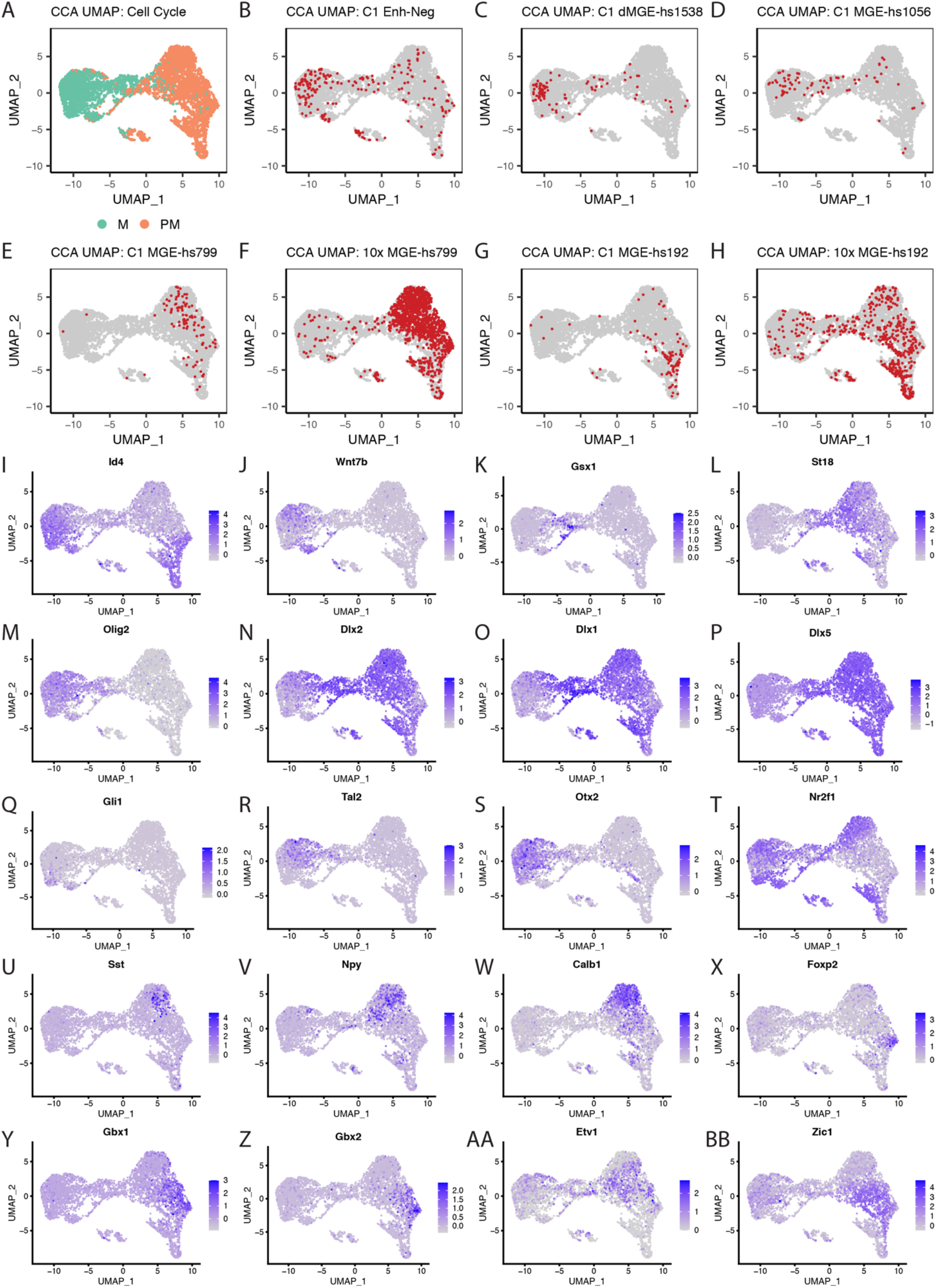
Canonical correlation analysis between C1 and 10x MGE scRNA-seq datasets. (**A**) TF-curated UMAP of combined C1 and 10x scRNA-seq using canonical correlation analysis, colored by mitotic state (green: mitotic; orange: postmitotic) (**B-H**) UMAP plots colored by C1 or 10x enhancer group. Red: selected group; grey: all other cells. (**I-BB**) UMAP plots of normalized gene expression for genes shown in Figure 5 (I-T) and Figure 6 (U-BB).

**Supplementary Figure 7:**
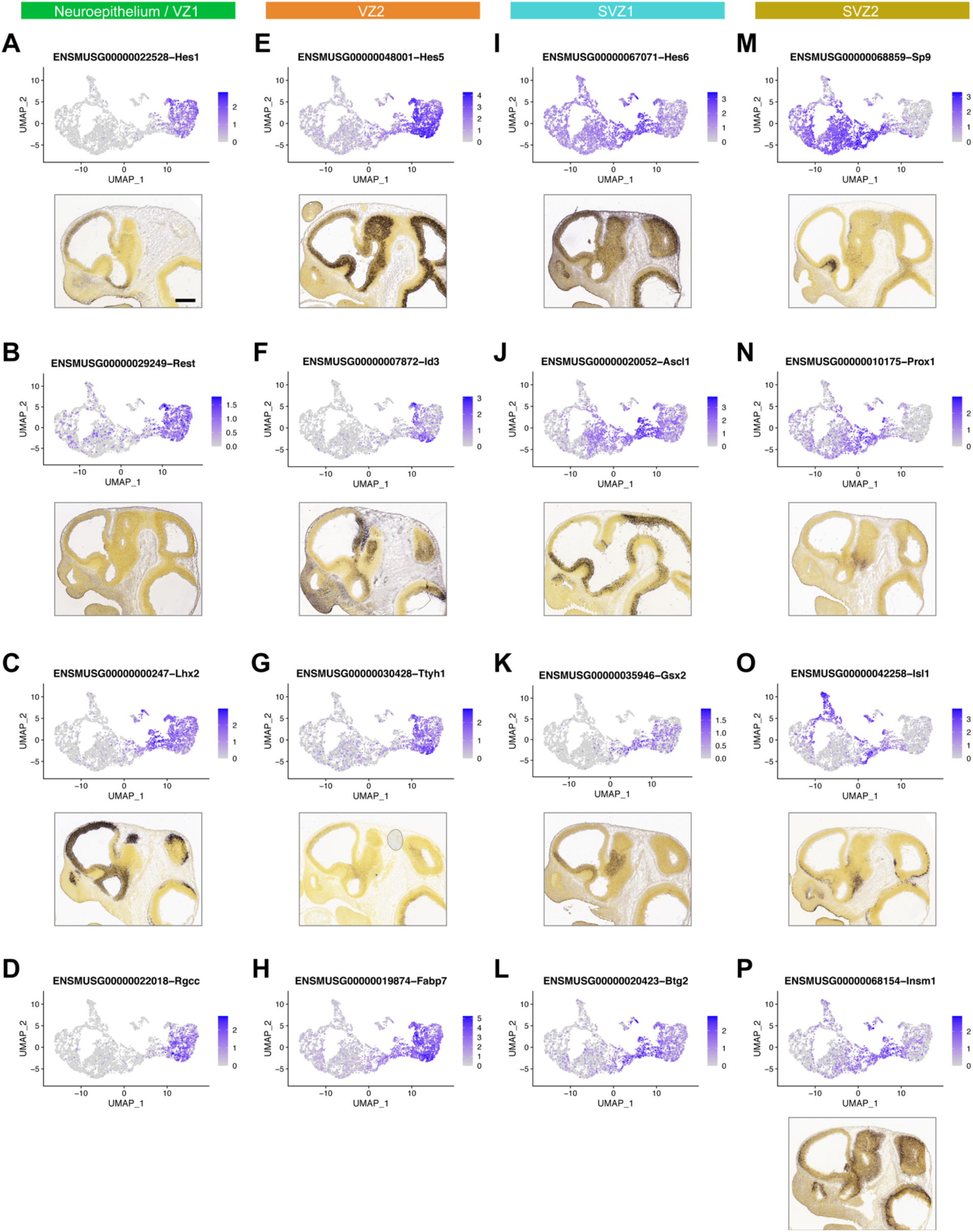
Mitotic genes marking VZ/SVZ transitions. (**A-F**) Gene expression UMAPs and representative ISH on E11.5 sagittal sections from the ABA showing expression of genes that correlate with developmental progression from early VZ1 (neuroepithelium) to late SVZ (SVZ2). Scale bar: 500 µm.

**Supplementary Figure 8:**
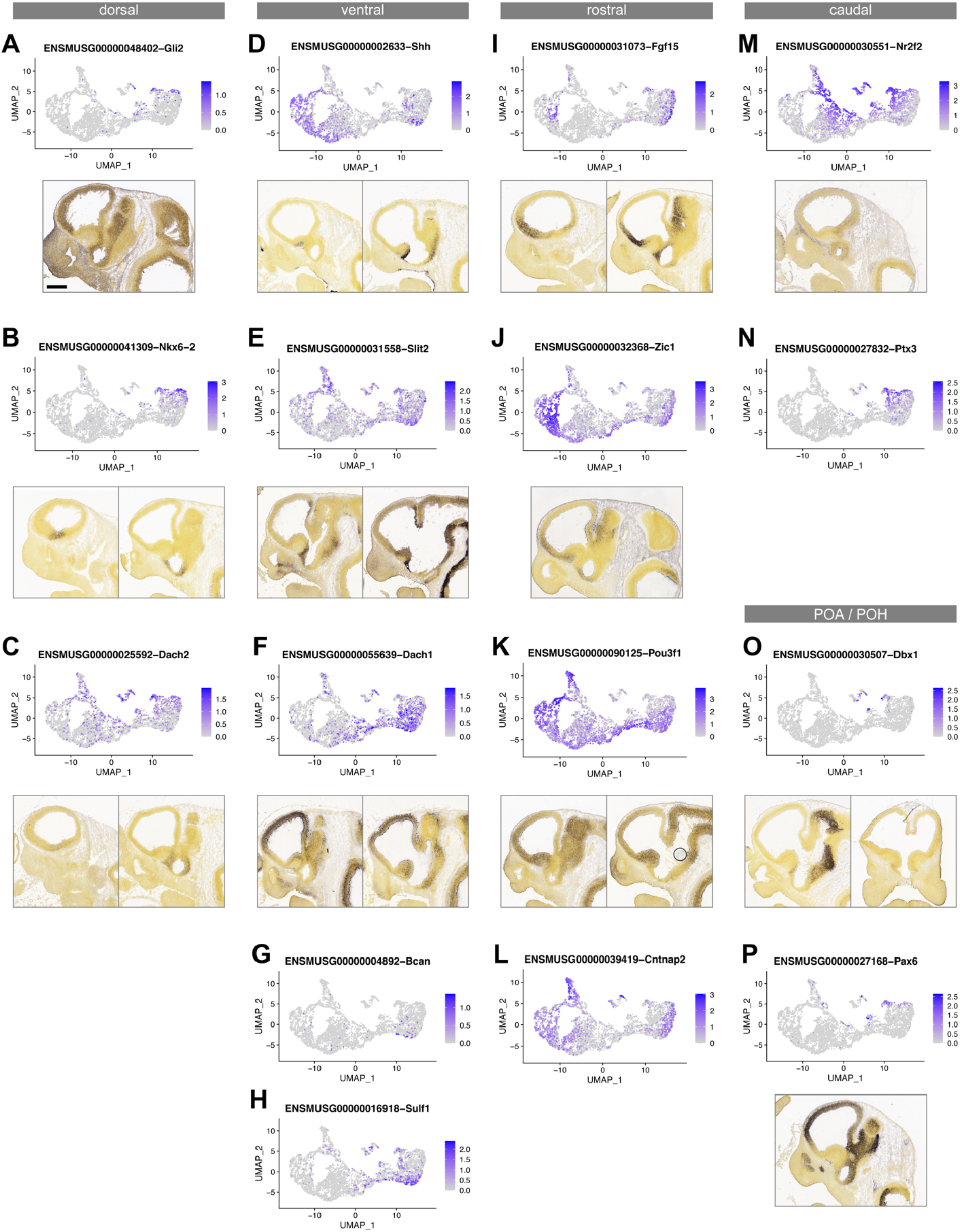
Mitotic genes with dorsoventral or rostrocaudal bias. (**A-H**) Gene expression UMAPs and representative ISH on E11.5 sagittal sections from the ABA showing genes with dorsal (*Gli2, Nkx6-2, Dach2*) or ventral (*Shh, Slit2, Dach1, Bcan, Sulf1*) bias in mitotic cells. *Dach2*, *Dach1*, *Bcan* and *Sulf1* regional expression in the MGE has not been reported previously. (**I-N**) Gene expression UMAPs and representative ISH on E11.5 sagittal sections from the ABA showing genes with rostrocaudal bias in mitotic cells (*Nr2f2, Ptx3, Fgf15, Cntnap2, Zic1, Pou3f1*). (**O**,**P**) UMAP plots and ISH images of genes marking POA2 (*Dbx1*) or POH (*Pax6*) VZ cells. These cells are contiguous with the caudal MGE anatomically and in the UMAP. Scale bar: 500 µm.

**Supplementary Figure 9:**
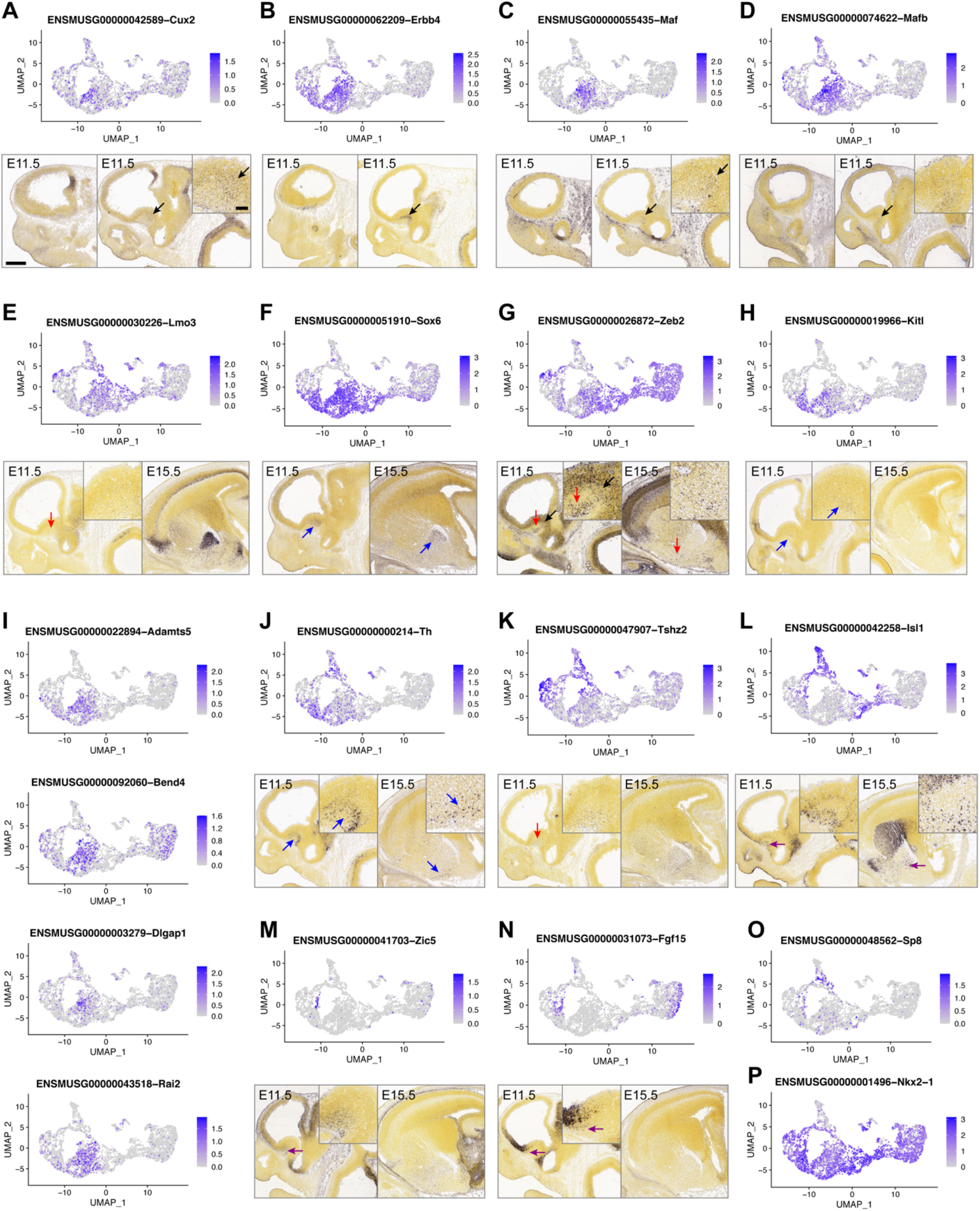
Genes marking classes of postmitotic neurons. (**A-D**) Gene expression UMAPs and representative ISH on E11.5 sagittal sections from the ABA showing expression of markers for early CINs (*Cux2, Erbb4, Maf, Mafb*). (**E-H**) Gene expression UMAPs and representative ISH for genes marking CINs and classes of projection neurons (*Lmo3, Sox6, Zeb2, Kitl*). (**I**) Gene expression UMAPs of proposed novel early CIN markers (*Adamts5, Bend4, Dlgap1, Rai2*). (**J-N**) Gene expression UMAPs and representative ISH for genes marking classes of MGE-derived projection neurons (*Th, Tshz2*) or cholinergic neurons (*Isl1, Zic5, Fgf15*). (**O**,**P**) UMAPs of *Sp8* and *Nkx2-1* showing the separation of cl-7 into an *Sp8*-positive non-MGE zone and an *Nkx2-1*-positive MGE-derived zone. Arrows indicate regions of higher magnification insets and cells of interest. Arrow colors: black, CINs; red, VP; blue, GP; purple; Ch. Scale bars: low magnification, 500 µm; high magnification insets, 100 µm.

**Supplementary Table 6:**
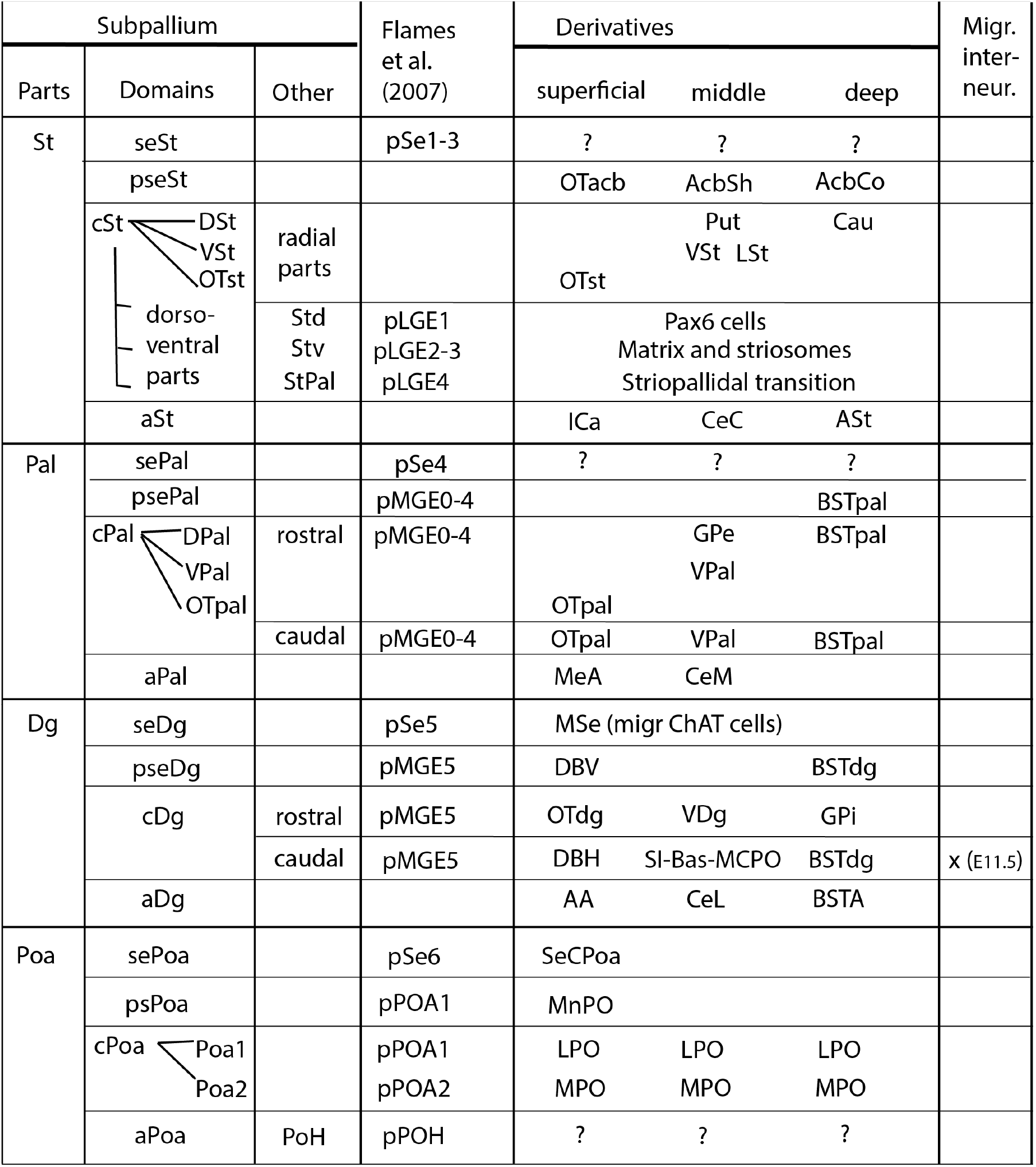
Proposed origins of subpallial neurons. Proposed origins of subpallial neurons, based on a body of anatomical, developmental and genoarchitectural findings (Puelles et al., 2016, 2013; Silberberg et al., 2016). Here we compared the nomenclature from this paper with that from (Flames et al., 2007). The left 3 columns (Subpallium) list the progenitor domains and subdomains: Column 1) Parts (major subdivisions): Striatum (St), Pallidum (Pal), Diagonal (Dg) and Preoptic Area (POA). Column 2) Domains (shared pattern across all the parts of septal, paraseptal, central and amygdalar regions, distinguished along the septoamygdalar axis): For St [septal St (seSt), paraseptal St (pseSt), central St (cSt), and amygdalar St (aSt)], for Pal [septal Pal (sePal), paraseptal Pal (psePal), central Pal (cPal), amygdalar Pal (aPal)], for Dg [septal Dg (seDg), paraseptal Dg (pseDg), central Dg (cDg), and amygdalar Dg (aDg)], and for Poa [septal Poa (sePoa), paraseptal Poa (psePoa), central Poa (cPoa), and amygdalar Poa (aPoa)]. Column 3) Other: provides morphological information on alternative subdivisions distinguished either along the radial dimension or the dorsoventral dimension. The central column lists the corresponding progenitor domains proposed in (Flames et al., 2007). Right columns (Derivatives) list the known neuronal derivatives of the subpallial progenitor domains, organized into radial (laminar) positions: superficial (closest to the pia), middle and deep (closest to the ventricle). The column on the far right (migrating interneurons) lists data from this publication proposing that the caudal Dg is the source for interneurons at E11.5. Other publications provide evidence that other subpallial progenitors also generate interneurons (for the pallium, striatum and olfactory bulb) at later ages. Other abbreviations, in the order that they are listed in the Table: OTacb (accumbens part of the olfactory tubercle), AcbSh (shell of the accumbens), AcbCo (core of the accumbens), OTst (central striatal part of the olfactory tubercle), Put and Cau (Putamen and Caudate, derivatives of the dorsal striatum, DSt), Vst, LSt (Ventral Striatum and Lateral Striatal Stripe, derivatives of the Ventral Striatum), Ica (intercalated nuclei of the amygdala), CeC (capsular part of the central amygdala), ASt (amygdalo-striatal transitional area), BSTpal (pallidal part of bed nucleus stria terminalis), GPe (external globus pallidus), VPal (ventral pallidum), OTpal (pallidal part of olfactory tubercle), MeA (medial amygdala), CeM (medial part of the central amygdala), MSe (medial septum, with cholinergic ChaAT neurons that may have migrated in from another region), DBV (diagonal band, vertical limb), BSTdg (diagonal part of bed nucleus stria terminalis), OTdg (diagonal part of olfactory tubercle), VDg (ventral diagonal area, GPi (internal globus pallidus), DBH (diagonal band, horizontal limb), SI-Bas-MCPO (substantia innominata, nucleus basalis, magnocellular preoptic complex), AA (anterior amygdala), CeL (lateral part of the central amygdala), BSTA (amygdalar part of BST), SeCPOA (septocommissural POA), MnPO (median preoptic nucleus), LPO (lateral preoptic area), MPO (medial preoptic area).

## References

Alexa, A., Rahnenfuhrer, J., 2020. topGO: Enrichment Analysis for Gene Ontology.

Anderson, S.A., Eisenstat, D.D., Shi, L., Rubenstein, J.L.R., 1997. Interneuron Migration from Basal Forebrain to Neocortex: Dependence on Dlx Genes. Science 278, 474–476. https://doi.org/10.1126/science.278.5337.474

Andrews, S., 2010. FastQC: a quality control tool for high throughput sequence data.

Angerer, P., Haghverdi, L., Büttner, M., Theis, F.J., Marr, C., Buettner, F., 2016. destiny: diffusion maps for large-scale single-cell data in R. Bioinformatics 32, 1241–1243. https://doi.org/10.1093/bioinformatics/btv715

Armoskus, C., Moreira, D., Bollinger, K., Jimenez, O., Taniguchi, S., Tsai, H.-W., 2014. Identification of sexually dimorphic genes in the neonatal mouse cortex and hippocampus. Brain Res. 1562, 23–38. https://doi.org/10.1016/j.brainres.2014.03.017

Asbreuk, C.H.J., van Schaick, H.S.A., Cox, J.J., Kromkamp, M., Smidt, M.P., Burbach, J.P.H., 2002. The homeobox genes Lhx7 and Gbx1 are expressed in the basal forebrain cholinergic system. Neuroscience 109, 287–298.

Batista-Brito, R., Ward, C., Fishell, G., 2020. The generation of cortical interneurons. Comprehensive Developmental Neuroscience. Patterning Cell Type Specif. Dev. CNS PNS 1, 461–480.

Beccari, L., Marco-Ferreres, R., Bovolenta, P., 2013. The logic of gene regulatory networks in early vertebrate forebrain patterning. Mech. Dev. 130, 95–111. https://doi.org/10.1016/j.mod.2012.10.004

Bedford, L., Walker, R., Kondo, T., van Crüchten, I., King, E.R., Sablitzky, F., 2005. Id4 is required for the correct timing of neural differentiation. Dev. Biol. 280, 386–395. https://doi.org/10.1016/j.ydbio.2005.02.001

Borello, U., Cobos, I., Long, J.E., Murre, C., Rubenstein, J.L., 2008. FGF15 promotes neurogenesis and opposes FGF8 function during neocortical development. Neural Develop. 3, 17. https://doi.org/10.1186/1749-8104-3-17

Breiman, L., 2001. Random Forests. Mach. Learn. 45, 5–32. https://doi.org/10.1023/A:1010933404324

Butler, A., Hoffman, P., Smibert, P., Papalexi, E., Satija, R., 2018. Integrating single-cell transcriptomic data across different conditions, technologies, and species. Nat. Biotechnol. 36, 411–420. https://doi.org/10.1038/nbt.4096

Campbell, P., Reep, R.L., Stoll, M.L., Ophir, A.G., Phelps, S.M., 2009. Conservation and diversity of Foxp2 expression in muroid rodents: Functional implications. J. Comp. Neurol. 512, 84–100. https://doi.org/10.1002/cne.21881

Carlson, M., 2019. org.Mm.eg.db: Genome wide annotation for Mouse.

Chen, L., Chatterjee, M., Li, J.Y.H., 2010. The mouse homeobox gene Gbx2 is required for the development of cholinergic interneurons in the striatum. J. Neurosci. Off. J. Soc. Neurosci. 30, 14824–14834.

Dobin, A., 2013. STAR: ultrafast universal RNA-seq aligner. Bioinformatics 29, 15–21.

Dunham, I., Kundaje, A., Aldred, S.F., Collins, P.J., Davis, C.A., Doyle, F., Epstein, C.B., Frietze, S., Harrow, J., Kaul, R., Khatun, J., Lajoie, B.R., Landt, S.G., Lee, B.-K., Pauli, F., Rosenbloom, K.R., Sabo, P., Safi, A., Sanyal, A., Shoresh, N., Simon, J.M., Song, L., Trinklein, N.D., Altshuler, R.C., Birney, E., Brown, J.B., Cheng, C., Djebali, S., Dong, X., Dunham, I., Ernst, J., Furey, T.S., Gerstein, M., Giardine, B., Greven, M., Hardison, R.C., Harris, R.S., Herrero, J., Hoffman, M.M., Iyer, S., Kellis, M., Khatun, J., Kheradpour, P., Kundaje, A., Lassmann, T., Li, Q., Lin, X., Marinov, G.K., Merkel, A., Mortazavi, A., Parker, S.C.J., Reddy, T.E., Rozowsky, J., Schlesinger, F., Thurman, R.E., Wang, J., Ward, L.D., Whitfield, T.W., Wilder, S.P., Wu, W., Xi, H.S., Yip, K.Y., Zhuang, J., Bernstein, B.E., Birney, E., Dunham, I., Green, E.D., Gunter, C., Snyder, M., Pazin, M.J., Lowdon, R.F., Dillon, L.A.L., Adams, L.B., Kelly, C.J., Zhang, J., Wexler, J.R., Green, E.D., Good, P.J., Feingold, E.A., Bernstein, B.E., Birney, E., Crawford, G.E., Dekker, J., Elnitski, L., Farnham, P.J., Gerstein, M., Giddings, M.C., Gingeras, T.R., Green, E.D., Guigó, R., Hardison, R.C., Hubbard, T.J., Kellis, M., Kent, W.J., Lieb, J.D., Margulies, E.H., Myers, R.M., Snyder, M., Stamatoyannopoulos, J.A., Tenenbaum, S.A., Weng, Z., White, K.P., Wold, B., Khatun, J., Yu, Y., Wrobel, J., Risk, B.A., Gunawardena, H.P., Kuiper, H.C., Maier, C.W., Xie, L., Chen, X., Giddings, M.C., Bernstein, B.E., Epstein, C.B., Shoresh, N., Ernst, J., Kheradpour, P., Mikkelsen, T.S., Gillespie, S., Goren, A., Ram, O., Zhang, X., Wang, L., Issner, R., Coyne, M.J., Durham, T., Ku, M., Truong, T., Ward, L.D., Altshuler, R.C., Eaton, M.L., Kellis, M., Djebali, S., Davis, C.A., Merkel, A., Dobin, A., Lassmann, T., Mortazavi, A., Tanzer, A., Lagarde, J., Lin, W., Schlesinger, F., Xue, C., Marinov, G.K., Khatun, J., Williams, B.A., Zaleski, C., Rozowsky, J., Röder, M., Kokocinski, F., Abdelhamid, R.F., Alioto, T., Antoshechkin, I., Baer, M.T., Batut, P., Bell, I., Bell, K., Chakrabortty, S., Chen, X., Chrast, J., Curado, J., Derrien, T., Drenkow, J., Dumais, E., Dumais, J., Duttagupta, R., Fastuca, M., Fejes-Toth, K., Ferreira, P., Foissac, S., Fullwood, M.J., Gao, H., Gonzalez, D., Gordon, A., Gunawardena, H.P., Howald, C., Jha, S., Johnson, R., Kapranov, P., King, B., Kingswood, C., Li, G., Luo, O.J., Park, E., Preall, J.B., Presaud, K., Ribeca, P., Risk, B.A., Robyr, D., Ruan, X., Sammeth, M., Sandhu, K.S., Schaeffer, L., See, L.-H., Shahab, A., Skancke, J., Suzuki, A.M., Takahashi, H., Tilgner, H., Trout, D., Walters, N., Wang, H., Wrobel, J., Yu, Y., Hayashizaki, Y., Harrow, J., Gerstein, M., Hubbard, T.J., Reymond, A., Antonarakis, S.E., Hannon, G.J., Giddings, M.C., Ruan, Y., Wold, B., Carninci, P., Guigó, R., Gingeras, T.R., Rosenbloom, K.R., Sloan, C.A., Learned, K., Malladi, V.S., Wong, M.C., Barber, G.P., Cline, M.S., Dreszer, T.R., Heitner, S.G., Karolchik, D., Kent, W.J., Kirkup, V.M., Meyer, L.R., Long, J.C., Maddren, M., Raney, B.J., Furey, T.S., Song, L., Grasfeder, L.L., Giresi, P.G., Lee, B.-K., Battenhouse, A., Sheffield, N.C., Simon, J.M., Showers, K.A., Safi, A., London, D., Bhinge, A.A., Shestak, C., Schaner, M.R., Ki Kim, S., Zhang, Z.Z., Mieczkowski, P.A., Mieczkowska, J.O., Liu, Z., McDaniell, R.M., Ni, Y., Rashid, N.U., Kim, M.J., Adar, S., Zhang, Z., Wang, T., Winter, D., Keefe, D., Birney, E., Iyer, V.R., Lieb, J.D., Crawford, G.E., Li, G., Sandhu, K.S., Zheng, M., Wang, P., Luo, O.J., Shahab, A., Fullwood, M.J., Ruan, X., Ruan, Y., Myers, R.M., Pauli, F., Williams, B.A., Gertz, J., Marinov, G.K., Reddy, T.E., Vielmetter, J., Partridge, E., Trout, D., Varley, K.E., Gasper, C., The ENCODE Project Consortium, Overall coordination (data analysis coordination), Data production leads (data production), Lead analysts (data analysis), Writing group, NHGRI project management (scientific management), Principal investigators (steering committee), Boise State University and University of North Carolina at Chapel Hill Proteomics groups (data production and analysis), Broad Institute Group (data production and analysis), Cold Spring Harbor, U. of G., Center for Genomic Regulation, Barcelona, RIKEN, Sanger Institute, University of Lausanne, Genome Institute of Singapore group (data production and analysis), Data coordination center at UC Santa Cruz (production data coordination), Duke University, E., University of Texas, Austin, University of North Carolina-Chapel Hill group (data production and analysis), Genome Institute of Singapore group (data production and analysis), HudsonAlpha Institute, C., UC Irvine, Stanford group (data production and analysis), 2012. An integrated encyclopedia of DNA elements in the human genome. Nature 489, 57–74. https://doi.org/10.1038/nature11247

Eisenstat, D.D., Liu, J.K., Mione, M., Zhong, W., Yu, G., Anderson, S.A., Ghattas, I., Puelles, L., Rubenstein, J.L.R., 1999. DLX-1, DLX-2, and DLX-5 expression define distinct stages of basal forebrain differentiation. J. Comp. Neurol. 414, 217–237. https://doi.org/10.1002/(SICI)1096-9861(19991115)414:2<217::AID-CNE6>3.0.CO;2-I

Elshatory, Y., Gan, L., 2008. The LIM-homeobox gene Islet-1 is required for the development of restricted forebrain cholinergic neurons. J. Neurosci. Off. J. Soc. Neurosci. 28, 3291–3297.

Feng, L., Hatten, M.E., Heintz, N., 1994. Brain lipid-binding protein (BLBP): A novel signaling system in the developing mammalian CNS. Neuron 12, 895–908. https://doi.org/10.1016/0896-6273(94)90341-7

Flames, N., Pla, R., Gelman, D.M., Rubenstein, J.L.R., Puelles, L., Marín, O., 2007. Delineation of multiple subpallial progenitor domains by the combinatorial expression of transcriptional codes. J. Neurosci. Off. J. Soc. Neurosci. 27, 9682–9695.

Flandin, P., Kimura, S., Rubenstein, J.L.R., 2010. The progenitor zone of the ventral medial ganglionic eminence requires Nkx2-1 to generate most of the globus pallidus but few neocortical interneurons. J. Neurosci. Off. J. Soc. Neurosci. 30, 2812–2823.

Fragkouli, A., Wijk, N.V. van, Lopes, R., Kessaris, N., Pachnis, V., 2009. LIM homeodomain transcription factor-dependent specification of bipotential MGE progenitors into cholinergic and GABAergic striatal interneurons. Development 136, 3841–3851. https://doi.org/10.1242/dev.038083

Hafemeister, C., Satija, R., 2019. Normalization and variance stabilization of single-cell RNA-seq data using regularized negative binomial regression. Genome Biol. 20, 296. https://doi.org/10.1186/s13059-019-1874-1

Haubensak, W., Attardo, A., Denk, W., Huttner, W.B., 2004. Neurons arise in the basal neuroepithelium of the early mammalian telencephalon: A major site of neurogenesis. Proc. Natl. Acad. Sci. 101, 3196–3201. https://doi.org/10.1073/pnas.0308600100

Hoch, R.V., Clarke, J.A., Rubenstein, J.L.R., 2015a. Fgf signaling controls the telencephalic distribution of Fgf-expressing progenitors generated in the rostral patterning center. Neural Develop. 10, 8–15.

Hoch, R.V., Lindtner, S., Price, J.D., Rubenstein, J.L.R., 2015b. OTX2 Transcription Factor Controls Regional Patterning within the Medial Ganglionic Eminence and Regional Identity of the Septum. Cell Rep. 12, 482–494.

Hu, J.S., Vogt, D., Lindtner, S., Sandberg, M., Silberberg, S.N., Rubenstein, J.L.R., 2017. Coup-TF1 and Coup-TF2 control subtype and laminar identity of MGE-derived neocortical interneurons. Dev. Camb. Engl. 144, 2837–2851.

Inoue, T., Ota, M., Ogawa, M., Mikoshiba, K., Aruga, J., 2007. Zic1 and Zic3 Regulate Medial Forebrain Development through Expansion of Neuronal Progenitors. J. Neurosci. 27, 5461–5473. https://doi.org/10.1523/JNEUROSCI.4046-06.2007

J.L.R., R., Campbell, K., 2020. Neurogenesis in the basal ganglia. Comprehensive Developmental Neuroscience. Patterning Cell Type Specif. Dev. CNS PNS 1, 399–403.

Kageyama, R., Ohtsuka, T., Kobayashi, T., 2008. Roles of Hes genes in neural development. Dev. Growth Differ. 50 Suppl 1, S97–103.

Kessaris, N., Magno, L., Rubin, A.N., Oliveira, M.G., 2014. Genetic programs controlling cortical interneuron fate. Curr. Opin. Neurobiol., SI: Inhibition: Synapses, Neurons and Circuits 26, 79–87. https://doi.org/10.1016/j.conb.2013.12.012

Kolde, R., 2015. pheatmap: Pretty heatmaps.

Kowalczyk, M.S., Tirosh, I., Heckl, D., Rao, T.N., Dixit, A., Haas, B.J., Schneider, R.K., Wagers, A.J., Ebert, B.L., Regev, A., 2015. Single-cell RNA-seq reveals changes in cell cycle and differentiation programs upon aging of hematopoietic stem cells. Genome Res. 25, 1860–1872. https://doi.org/10.1101/gr.192237.115

Lein, E.S., Hawrylycz, M.J., Ao, N., Ayres, M., Bensinger, A., Bernard, A., Boe, A.F., Boguski, M.S., Brockway, K.S., Byrnes, E.J., Chen, Lin, Chen, Li, Chen, T.-M., Chi Chin, M., Chong, J., Crook, B.E., Czaplinska, A., Dang, C.N., Datta, S., Dee, N.R., Desaki, A.L., Desta, T., Diep, E., Dolbeare, T.A., Donelan, M.J., Dong, H.-W., Dougherty, J.G., Duncan, B.J., Ebbert, A.J., Eichele, G., Estin, L.K., Faber, C., Facer, B.A., Fields, R., Fischer, S.R., Fliss, T.P., Frensley, C., Gates, S.N., Glattfelder, K.J., Halverson, K.R., Hart, M.R., Hohmann, J.G., Howell, M.P., Jeung, D.P., Johnson, R.A., Karr, P.T., Kawal, R., Kidney, J.M., Knapik, R.H., Kuan, C.L., Lake, J.H., Laramee, A.R., Larsen, K.D., Lau, C., Lemon, T.A., Liang, A.J., Liu, Y., Luong, L.T., Michaels, J., Morgan, J.J., Morgan, R.J., Mortrud, M.T., Mosqueda, N.F., Ng, L.L., Ng, R., Orta, G.J., Overly, C.C., Pak, T.H., Parry, S.E., Pathak, S.D., Pearson, O.C., Puchalski, R.B., Riley, Z.L., Rockett, H.R., Rowland, S.A., Royall, J.J., Ruiz, M.J., Sarno, N.R., Schaffnit, K., Shapovalova, N.V., Sivisay, T., Slaughterbeck, C.R., Smith, S.C., Smith, K.A., Smith, B.I., Sodt, A.J., Stewart, N.N., Stumpf, K.-R., Sunkin, S.M., Sutram, M., Tam, A., Teemer, C.D., Thaller, C., Thompson, C.L., Varnam, L.R., Visel, A., Whitlock, R.M., Wohnoutka, P.E., Wolkey, C.K., Wong, V.Y., Wood, M., Yaylaoglu, M.B., Young, R.C., Youngstrom, B.L., Feng Yuan, X., Zhang, B., Zwingman, T.A., Jones, A.R., 2007. Genome-wide atlas of gene expression in the adult mouse brain. Nature 445, 168–176. https://doi.org/10.1038/nature05453

Levine, M., 2008. A systems view of Drosophila segmentation. Genome Biol. 9, 207. https://doi.org/10.1186/gb-2008-9-2-207

Liao, Y., Smyth, G.K., Shi, W., 2014. featureCounts: an efficient general purpose program for assigning sequence reads to genomic features. Bioinformatics 30, 923–930.

Lim, L., Mi, D., Llorca, A., Marín, O., 2018. Development and Functional Diversification of Cortical Interneurons. Neuron 100, 294–313. https://doi.org/10.1016/j.neuron.2018.10.009

Lindtner, S., Catta-Preta, R., Tian, H., Su-Feher, L., Price, J.D., Dickel, D.E., Greiner, V., Silberberg, S.N., McKinsey, G.L., McManus, M.T., Pennacchio, L.A., Visel, A., Nord, A.S., Rubenstein, J.L.R., 2019. Genomic Resolution of DLX-Orchestrated Transcriptional Circuits Driving Development of Forebrain GABAergic Neurons. Cell Rep. 28, 2048–2063.e8. https://doi.org/10.1016/j.celrep.2019.07.022

Long, J.E., Garel, S., Alvarez-Dolado, M., Yoshikawa, K., Osumi, N., Alvarez-Buylla, A., Rubenstein, J.L.R., 2007. Dlx-Dependent and -Independent Regulation of Olfactory Bulb Interneuron Differentiation. J. Neurosci. 27, 3230–3243. https://doi.org/10.1523/JNEUROSCI.5265-06.2007

Long, J.E., Swan, C., Liang, W.S., Cobos, I., Potter, G.B., Rubenstein, J.L.R., 2009. Dlx1&2 and Mash1 transcription factors control striatal patterning and differentiation through parallel and overlapping pathways. J. Comp. Neurol. 512, 556–572.

Madisen, L., Zwingman, T.A., Sunkin, S.M., Oh, S.W., Zariwala, H.A., Gu, H., Ng, L.L., Palmiter, R.D., Hawrylycz, M.J., Jones, A.R., Lein, E.S., Zeng, H., 2010. A robust and high-throughput Cre reporting and characterization system for the whole mouse brain. Nat. Neurosci. 13, 133–140.

Magno, L., Barry, C., Schmidt-Hieber, C., Theodotou, P., Häusser, M., Kessaris, N., 2017. NKX2-1 Is Required in the Embryonic Septum for Cholinergic System Development, Learning, and Memory. Cell Rep. 20, 1572–1584.

Marín, O., Anderson, S.A., Rubenstein, J.L.R., 2000. Origin and Molecular Specification of Striatal Interneurons. J. Neurosci. 20, 6063–6076. https://doi.org/10.1523/JNEUROSCI.20-16-06063.2000

Mayer, C., Hafemeister, C., Bandler, R.C., Machold, R., Batista Brito, R., Jaglin, X., Allaway, K., Butler, A., Fishell, G., Satija, R., 2018. Developmental diversification of cortical inhibitory interneurons. Nature 555, 457–462. https://doi.org/10.1038/nature25999

McGinnis, C.S., Patterson, D.M., Winkler, J., Conrad, D.N., Hein, M.Y., Srivastava, V., Hu, J.L., Murrow, L.M., Weissman, J.S., Werb, Z., Chow, E.D., Gartner, Z.J., 2019. MULTI-seq: sample multiplexing for single-cell RNA sequencing using lipid-tagged indices. Nat. Methods 16, 619–626.

McGregor, M.M., McKinsey, G.L., Girasole, A.E., Bair-Marshall, C.J., Rubenstein, J.L.R., Nelson, A.B., 2019. Functionally Distinct Connectivity of Developmentally Targeted Striosome Neurons. Cell Rep. 29, 1419–1428.e5. https://doi.org/10.1016/j.celrep.2019.09.076

McInnes, L., Healy, J., Saul, N., Großberger, L., 2018. UMAP: Uniform Manifold Approximation and Projection. J. Open Source Softw. 3, 861. https://doi.org/10.21105/joss.00861

Mckinsey, G.L., Lindtner, S., Trzcinski, B., Visel, A., Pennacchio, L.A., Huylebroeck, D., Higashi, Y., Rubenstein, J.L.R., 2013. Dlx1&2-dependent expression of Zfhx1b (Sip1, Zeb2) regulates the fate switch between cortical and striatal interneurons. Neuron 77, 83–98.

Mi, D., Li, Z., Lim, L., Li, M., Moissidis, M., Yang, Y., Gao, T., Hu, T.X., Pratt, T., Price, D.J., Sestan, N., Marín, O., 2018. Early emergence of cortical interneuron diversity in the mouse embryo. Science 360, 81–85. https://doi.org/10.1126/science.aar6821

Nóbrega-Pereira, S., Gelman, D., Bartolini, G., Pla, R., Pierani, A., Marín, O., 2010. Origin and Molecular Specification of Globus Pallidus Neurons. J. Neurosci. 30, 2824–2834. https://doi.org/10.1523/JNEUROSCI.4023-09.2010

Nord, A.S., 2015. Learning about mammalian gene regulation from functional enhancer assays in the mouse. Genomics, Recent advances in functional assays of transcriptional enhancers 106, 178–184. https://doi.org/10.1016/j.ygeno.2015.06.008

Nord, A.S., 2013. Rapid and pervasive changes in genome-wide enhancer usage during mammalian development. Cell 155, 1521–1531.

Pai, E.L.-L., Vogt, D., Clemente-Perez, A., Mckinsey, G.L., Cho, F.S., Hu, J.S., Wimer, M., Paul, A., Fazel Darbandi, S., Pla, R., Nowakowski, T.J., Goodrich, L.V., Paz, J.T., Rubenstein, J.L.R., 2019. Mafb and c-Maf Have Prenatal Compensatory and Postnatal Antagonistic Roles in Cortical Interneuron Fate and Function. Cell Rep. 26, 1157–1173.e5.

Pattabiraman, K., Golonzhka, O., Lindtner, S., Nord, A.S., Taher, L., Hoch, R., Silberberg, S.N., Zhang, D., Chen, B., Zeng, H., Pennacchio, L.A., Puelles, L., Visel, A., Rubenstein, J.L.R., 2014. Transcriptional Regulation of Enhancers Active in Protodomains of the Developing Cerebral Cortex. Neuron 82, 989–1003. https://doi.org/10.1016/j.neuron.2014.04.014

Petryniak, M.A., Potter, G.B., Rowitch, D.H., Rubenstein, J.L.R., 2007. Dlx1 and Dlx2 control neuronal versus oligodendroglial cell fate acquisition in the developing forebrain. Neuron 55, 417–433.

Picard Toolkit, 2019. Broad Institute, GitHub Repository.

Porteus, M.H., Bulfone, A., Liu, J.K., Puelles, L., Lo, L.C., Rubenstein, J.L., 1994. DLX-2, MASH-1, and MAP-2 expression and bromodeoxyuridine incorporation define molecularly distinct cell populations in the embryonic mouse forebrain. J. Neurosci. 14, 6370–6383. https://doi.org/10.1523/JNEUROSCI.14-11-06370.1994

Preibisch, S., Saalfeld, S., Tomancak, P., 2009. Globally optimal stitching of tiled 3D microscopic image acquisitions. Bioinformatics 25, 1463–1465.

Puelles, L., Harrison, M., Paxinos, G., Watson, C., 2013. A developmental ontology for the mammalian brain based on the prosomeric model. Trends Neurosci. 36, 570–578. https://doi.org/10.1016/j.tins.2013.06.004

Puelles, L., Morales-Delgado, N., Merchán, P., Castro-Robles, B., Martínez-de-la-Torre, M., Díaz, C., Ferran, J.L., 2016. Radial and tangential migration of telencephalic somatostatin neurons originated from the mouse diagonal area. Brain Struct. Funct. 221, 3027–3065.

Reddington, J.P., Garfield, D.A., Sigalova, O.M., Karabacak Calviello, A., Marco-Ferreres, R., Girardot, C., Viales, R.R., Degner, J.F., Ohler, U., Furlong, E.E.M., 2020. Lineage-Resolved Enhancer and Promoter Usage during a Time Course of Embryogenesis. Dev. Cell 55, 648–664.e9. https://doi.org/10.1016/j.devcel.2020.10.009

Roychoudhury, K., Salomone, J., Qin, S., Cain, B., Adam, M., Potter, S.S., Nakafuku, M., Gebelein, B., Campbell, K., 2020. Physical interactions between Gsx2 and Ascl1 balance progenitor expansion versus neurogenesis in the mouse lateral ganglionic eminence. Dev. Camb. Engl. 147, dev185348.

Rubin, A.N., Alfonsi, F., Humphreys, M.P., Choi, C.K.P., Rocha, S.F., Kessaris, N., 2010. The Germinal Zones of the Basal Ganglia But Not the Septum Generate GABAergic Interneurons for the Cortex. J. Neurosci. 30, 12050–12062. https://doi.org/10.1523/JNEUROSCI.6178-09.2010

Sanchez-Ortiz, E., Yui, D., Song, D., Li, Y., Rubenstein, J.L., Reichardt, L.F., Parada, L.F., 2012. TrkA Gene Ablation in Basal Forebrain Results in Dysfunction of the Cholinergic Circuitry. J. Neurosci. 32, 4065–4079. https://doi.org/10.1523/JNEUROSCI.6314-11.2012

Sandberg, M., Flandin, P., Silberberg, S., Su-Feher, L., Price, J.D., Hu, J.S., Kim, C., Visel, A., Nord, A.S., Rubenstein, J.L.R., 2016. Transcriptional Networks Controlled by NKX2-1 in the Development of Forebrain GABAergic Neurons. Neuron 91, 1260–1275. https://doi.org/10.1016/j.neuron.2016.08.020

Silberberg, S.N., Taher, L., Lindtner, S., Sandberg, M., Nord, A.S., Vogt, D., Mckinsey, G.L., Hoch, R., Pattabiraman, K., Zhang, D., Ferran, J.L., Rajkovic, A., Golonzhka, O., Kim, C., Zeng, H., Puelles, L., Visel, A., Rubenstein, J.L.R., 2016. Subpallial Enhancer Transgenic Lines: a Data and Tool Resource to Study Transcriptional Regulation of GABAergic Cell Fate. Neuron 92, 59–74. https://doi.org/10.1016/j.neuron.2016.09.027

Stuart, T., Butler, A., Hoffman, P., Hafemeister, C., Papalexi, E., Mauck, W.M., Hao, Y., Stoeckius, M., Smibert, P., Satija, R., 2019. Comprehensive Integration of Single-Cell Data. Cell 177, 1888–1902.e21. https://doi.org/10.1016/j.cell.2019.05.031

Visel, A., Taher, L., Girgis, H., May, D., Golonzhka, O., Hoch, R.V., McKinsey, G.L., Pattabiraman, K., Silberberg, S.N., Blow, M.J., Hansen, D.V., Nord, A.S., Akiyama, J.A., Holt, A., Hosseini, R., Phouanenavong, S., Plajzer-Frick, I., Shoukry, M., Afzal, V., Kaplan, T., Kriegstein, A.R., Rubin, E.M., Ovcharenko, I., Pennacchio, L.A., Rubenstein, J.L.R., 2013. A High-Resolution Enhancer Atlas of the Developing Telencephalon. Cell 152, 895–908. https://doi.org/10.1016/j.cell.2012.12.041

Wang, L., Wang, S., Li, W., 2012. RSeQC: quality control of RNA-seq experiments. Bioinformatics 28, 2184–2185.

Yun, K., Mantani, A., Garel, S., Rubenstein, J., Israel, M.A., 2004. Id4 regulates neural progenitor proliferation and differentiation in vivo. Development 131, 5441–5448. https://doi.org/10.1242/dev.01430

Yuzwa, S.A., Borrett, M.J., Innes, B.T., Voronova, A., Ketela, T., Kaplan, D.R., Bader, G.D., Miller, F.D., 2017. Developmental Emergence of Adult Neural Stem Cells as Revealed by Single-Cell Transcriptional Profiling. Cell Rep. 21, 3970–3986.

Zeisel, A., Hochgerner, H., Lönnerberg, P., Johnsson, A., Memic, F., van der Zwan, J., Häring, M., Braun, E., Borm, L.E., La Manno, G., Codeluppi, S., Furlan, A., Lee, K., Skene, N., Harris, K.D., Hjerling-Leffler, J., Arenas, E., Ernfors, P., Marklund, U., Linnarsson, S., 2018. Molecular Architecture of the Mouse Nervous System. Cell 174, 999–1014.e22. https://doi.org/10.1016/j.cell.2018.06.021

Zheng, G.X.Y., Terry, J.M., Belgrader, P., Ryvkin, P., Bent, Z.W., Wilson, R., Ziraldo, S.B., Wheeler, T.D., McDermott, G.P., Zhu, J., Gregory, M.T., Shuga, J., Montesclaros, L., Underwood, J.G., Masquelier, D.A., Nishimura, S.Y., Schnall-Levin, M., Wyatt, P.W., Hindson, C.M., Bharadwaj, R., Wong, A., Ness, K.D., Beppu, L.W., Deeg, H.J., McFarland, C., Loeb, K.R., Valente, W.J., Ericson, N.G., Stevens, E.A., Radich, J.P., Mikkelsen, T.S., Hindson, B.J., Bielas, J.H., 2017. Massively parallel digital transcriptional profiling of single cells. Nat. Commun. 8, 14049. https://doi.org/10.1038/ncomms14049

